# Deciphering sources of PET signals in the tumor microenvironment of glioblastoma at cellular resolution

**DOI:** 10.1101/2023.01.26.522174

**Authors:** Laura M Bartos, Sabrina V Kirchleitner, Zeynep Ilgin Kolabas, Stefanie Quach, Jens Blobner, Stephan A Mueller, Selin Ulukaya, Luciano Hoeher, Izabela Horvath, Karin Wind-Mark, Adrien Holzgreve, Viktoria C Ruf, Lukas Gold, Lea H Kunze, Sebastian T Kunte, Philipp Beumers, Melissa Antons, Artem Zatcepin, Nils Briel, Leonie Hoermann, Denise Messerer, Peter Bartenstein, Markus J Riemenschneider, Simon Lindner, Sibylle Ziegler, Jochen Herms, Stefan F Lichtenthaler, Ali Ertürk, Joerg C Tonn, Louisa von Baumgarten, Nathalie L Albert, Matthias Brendel

## Abstract

Various cellular sources hamper interpretation of positron-emission-tomography (PET) biomarkers in the tumor microenvironment (TME). We developed immunomagnetic cell sorting after *in vivo* radiotracer injection (scRadiotracing) in combination with 3D-histology via tissue clearing to dissect the cellular allocation of PET signals in the TME. In SB28 glioblastoma mice, translocator protein (TSPO) radiotracer uptake per tumor cell was higher compared to tumor-associated microglia/macrophages (TAMs). Cellular radiotracer uptake was validated by proteomics and confirmed for *in vitro* samples of patients with glioblastoma. Regional agreement between PET signals and single cell tracer uptake predicted the individual cell distribution in 3D-histology. In consideration of cellular tracer uptake and cell type abundance, tumor cells were the main contributor to TSPO enrichment in glioblastoma, however proteomics identified potential PET targets highly specific for TAMs. Combining cellular tracer uptake measures with 3D-histology facilitates precise allocation of complex PET signal sources and will serve to validate novel TAM-specific radioligands.

## Introduction

The tumor microenvironment (TME) emerged as an acknowledged key component in cancer biology and treatment^1^. In particular, novel immunotherapies against targets of the TME facilitate potentiation of host antitumor immune responses^2^. However, despite the need to monitor novel anti-cancer therapeutics *in vivo*, specific biomarkers for immune cells of the TME are still limited^3^. Recent efforts originated in radiolabeled antibodies for positron-emission-tomography (PET) that specifically target immune cells^4^, but the cell-type heterogeneity of the TME hampers cellular allocation and interpretation of such biomarkers. Furthermore, antibody-based PET biomarkers have limited penetrance through the blood-brain-barrier, which limits their use in brain malignancies, such as glioblastoma which represents the most common malignant primary brain tumor with a very poor prognosis^5^. In this disease, 18 kDa translocator protein (TSPO)-PET showed opportunities to assess myeloid cells of the TME as important contributors to immune evasion of glioblastoma^6^. However, the detailed sources of TSPO and other TME biomarkers in glioma have not yet been thoroughly elucidated, which again hampers interpretation and understanding of individual PET images. In glioma, tumor-associated microglia and macrophages (TAMs) show elevated expression of TSPO *in vitro*^7^ and *in vivo*^6^, acting as a potential signal target of TSPO ligands. However, this finding is a matter of debate since in high-grade tumors, TSPO was found to be highly overexpressed by tumor cells as well^8,9,10^.

To overcome the limitations of previous immunohistochemistry to PET comparisons, we aimed to resolve the complex sources of TME biomarkers in glioma at cellular resolution. We therefore established and validated a novel approach of immunomagnetic cell sorting after radiotracer injection (single cell Radiotracing, scRadiotracing)^11^ and exploited this technique in a mouse model of experimental glioblastoma. TSPO tracer uptake was measured in single tumor cells and CD11b-positive immune cells of a SB28 glioblastoma mouse model, validated against TSPO protein levels as determined by proteomics as a proof of concept. Furthermore, we investigated associations between single cell TSPO tracer uptake and tumor heterogeneity in TSPO-PET. To bridge the gap between implantation of murine tumors and spontaneous glioma development in human brain, we transferred TSPO scRadiotracing to human samples and determined proportions of TSPO tracer uptake *in vitro* in tumor and immune cells of patients with high-grade and low-grade glioma immediately after surgery. As a second major innovation, 3D-histology via tissue clearing and light sheet microscopy was integrated to acknowledge PET signals as a composite of cell type abundance and single cell tracer uptake. Finally, the full proteome of isolated SB28 tumor cells and TAMs was analyzed for identification of potential TAM-specific TME radioligand targets with superior binding capacity when compared to TSPO.

## Results

### Single cell tracer uptake measures of microglia as calculated by scRadiotracing correspond to PET signal changes upon microglia depletion

Immune PET biomarkers such as TSPO have the potential to monitor TAMs of the TME, but the target signal of currently available TSPO radioligands is challenged by several cellular sources. To validate scRadiotracing for the TSPO tracer [^18^F]GE-180, we compared the magnitude of the microglia-specific TSPO-PET signal with the recovered radioactivity in isolated microglia of untreated mice. First, we investigated the allocation of the TSPO tracer in the healthy mouse brain. Isolated CD11b(+) microglia (1.2 ± 0.1 × 10^5^ cells) of untreated mice revealed 7.7 ± 0.7 × 10^−7^ percentage radioactivity (normalized to injected dose (ID) and body weight (BW), %ID*BW) per single cell, whereas enriched ACSA2(+) astrocytes (5.2 ± 0.7 × 10^5^ cells) indicated 12.5-fold lower radioactivity per single cell (6.2 ± 0.7 × 10^−8^ %ID*BW; p = 0.00016; **Fig. 1A**), speaking for strong specificity of the tracer for microglia over astrocytes. Next, extrapolation by published microglia cell numbers (6% of all brain cells, 7.63 × 10^6^ microglia cells) from dedicated studies^12^, whole brain radioactivity (0.19 - 0.30 MBq) and %brain dose per single microglia cell (1.8 × 10^−6^ - 3.6 × 10^−6^) were used to calculate the contribution of the microglia population to the radioactivity in the whole brain to be 17.5% ± 2.2%.

**Figure 1:**
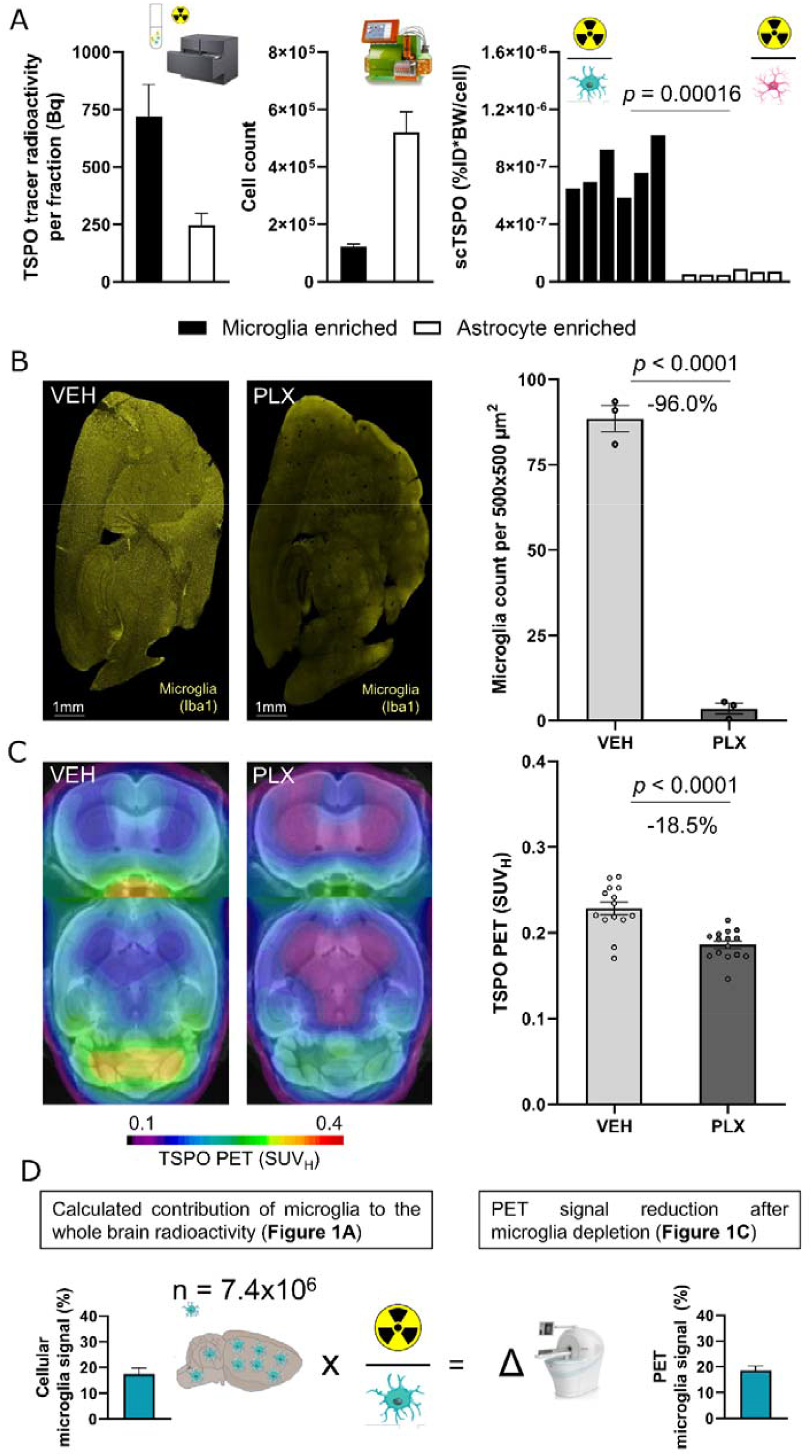
TSPO-PET signal reduction in the whole brain after microglia depletion corresponds to the whole brain signal attributable to tracer uptake in microglia. (**A**) Components of scRadiotracing after *in vivo* tracer injection. Upon *in vivo* tracer injection and magnetic cell separation, cell pellets were analyzed by a high sensitive gamma counter to measure the radioactivity (Bq = Becquerel) in the sample (left panel, n=6, mean±SEM) as well as by flow cytometry to determine the cell count (middle panel, n=6, mean±SEM). After calculation of radioactivity per single cell (scTSPO, normalized to injected dose (ID) and body weight (BW)) in each sample, untreated mice showed significantly higher TSPO tracer uptake in microglia when compared to astrocytes (right panel; each bar represents a single animal; n=6, paired t-test). (**C**) Representative sagittal Iba1 immunohistochemistry sections of VEH (n=3) and PLX5622 treated (n=3) mice and quantification of stained Iba1 area (%). Unpaired t-test, mean±SEM. (**B**) Group average TSPO-PET images of healthy mice upon a MRI template after vehicle treatment (VEH, n=14) or CSF1R inhibition (PLX5622, microglia depletion, n=15) indicated a distinct signal reduction in the whole brain of PLX treated animals (−18.5%). SUV_H_ = heart normalized standardized uptake value (SUV). Unpaired t-test, mean±SEM. (**D**) Extrapolation of the signal attributable to microglia using published cell numbers^43^ and tracer uptake per single microglia cell as calculated by scRadiotracing. The calculated 17.5% contribution of microglia to the whole brain TSPO-PET signal corresponded to the observed TSPO-PET signal reduction after microglia depletion (18.5%). mean±SEM.

To quantify the contribution of microglia to the overall TSPO-PET signal, we used CSF1R inhibition via PLX5622 and hereby depleted 96% of all microglia in the mouse brain (**Fig. 1B**). As a result of microglia depletion, the TSPO-PET signal (**Fig. 1C**) in mice at 10.1 ± 2.1 months of age^13,14^ showed 18.5% lower standardized uptake values (SUV_H_) when compared to vehicle controls (0.19 ± 0.01 vs. 0.23 ± 0.01; p < 0.0001). This decrease corresponded to the contribution of the microglia cell population extrapolated after scRadiotracing (p = 0.770; **Fig. 1D**). Thus, TSPO scRadiotracing provided a nearly complete recovery of the microglia-specific PET signal in healthy mice and suggested capability of the method to be used for tracer allocation in cells of the TME.

### TSPO tracer uptake of tumor cells and TAMs reflects cellular TSPO protein levels in the SB28 glioblastoma mouse model

In the next step, we applied scRadiotracing to an experimental orthotopic glioblastoma model (SB28) that closely reflects the human TME^15^ to investigate the contribution of tumor cells and TAMs to the TSPO-PET signal (**Fig. 2A-C**). Using MACS, CD11b(+) immune cells were isolated from tumor and sham injected brains (TAMs or sham microglia), tumor samples underwent subsequent tumor cell isolation. Residual cell pellets were analyzed as depleted fractions. We extracted 1.4 ± 0.4 × 10^5^ GFP(+) tumor cells and 6.7 ± 1.7 × 10^5^ CD11b(+) TAMs from tumor mice and 4.2 ± 0.8 × 10^4^ CD11b(+) microglia from sham mice (**Fig. 2A**) in enriched fractions at high purity as determined by flow cytometry (tumor cells: 87% ± 2%; TAMs: 90% ± 1%; sham microglia: 89% ± 1%; **Fig. 2B-D**). Signal-to-noise-ratios (SNRs) for radioactivity detection were consistently ≥ 2 (**Fig. 2A**).

**Figure 2:**
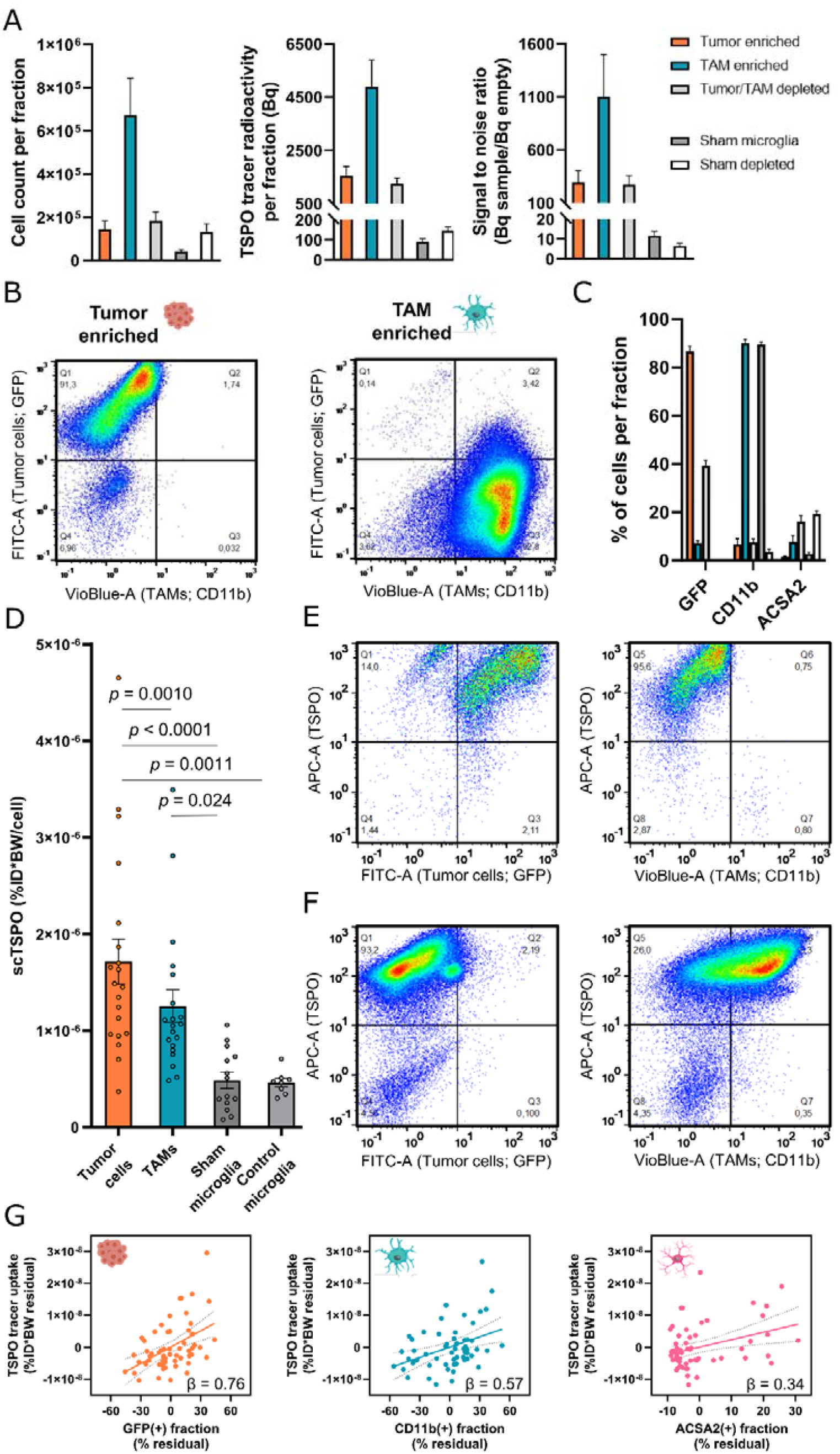
scRadiotracing in the SB28 glioblastoma mouse model allows differential assessment of TSPO tracer uptake in tumor cells and TAMs. **(A)** Acquired components of scRadiotracing after *in vivo* tracer injection in SB28 glioblastoma (n=20) and sham (n=14) mice. Using immunomagnetic cell sorting, tumor cells (GFP(+)) and CD11b(+) TAMs from SB28 bearing mice and CD11b(+) microglia from sham injected mice were enriched, thus leaving the respective residual depleted cell fractions. Absolute cell numbers (left panel) and measured radioactivity (middle panel) resulted in satisfying signal to noise ratios (right panel) for all enriched and depleted fractions investigated. mean±SEM. **(B)** Immunomagnetic cell sorting of tumor cells and TAMs led to >90% purity in enriched fractions as confirmed by flow cytometry. **(c)** Distribution of tumor cells (GFP), TAMs (CD11b) and astrocytes (ACSA2) in enriched and depleted fractions of SB28 glioblastoma (n=20) and sham (n=14) mice. High purity was reached for tumor and TAM enriched fractions, while remaining depleted fractions still contained tumor cells but no TAMs. Astrocytes could be detected in both depleted fractions using ACSA2. mean±SEM. **(D)** Comparison of the single cell TSPO tracer uptake of isolated tumor cells (n=20) and TAMs (n=20) in SB28 glioblastoma mice as well as microglia of sham (n=14) and untreated control (n=8) mice. Tumor cells and TAMs showed significantly higher TSPO tracer uptake per single cell (scTSPO, normalized to injected dose (ID) and body weight (BW)) when compared to isolated microglia of sham and untreated control animals. In SB28 mice, tumor cells showed higher single cell TSPO tracer uptake when compared to TAMs. Single cell tracer uptake of sham microglia showed no difference when compared to control microglia of untreated mice. Paired t-test for tumor cells vs. TAMs, one-way ANOVA for all other comparisons, mean±SEM. (**E**,**F**) TSPO co-staining in flow-cytometry shows that nearly all tumor cells (GFP; **E**) and TAMs (CD11b; **F**) were also positive for TSPO. Notably, the minor population of CD11b(−) cells in the TAM enriched fraction did not show positivity for TSPO (**F**, lower left quadrant), confirming the specificity of the TSPO co-staining. Pooled data from n=3 tumors. (**G**) Regression model including the enriched and depleted fractions of SB28 mice indicated highest contribution of tumor cells (left) to the radioactivity in the sample, followed by TAMs (middle) and astrocytes (right). Linear regression, β = standardized regression coefficient, n=60 samples, error bands represent 95% confidence interval.

Subsequently, single cell tracer uptake of tumor cells and TAMs was quantified to decipher cellular contributions to the TSPO tracer uptake in glioblastoma. Both, single SB28 tumor cells (3.53-fold, 1.7 ± 0.2 × 10^−6^ %ID*BW; p < 0.0001, 1-way ANOVA; **Fig. 2D**) and TAMs (2.58-fold, 1.3 ± 0.2 × 10^−6^ %ID*BW; p = 0.024, 1-way ANOVA; **Fig. 2D**) indicated higher TSPO tracer uptake when compared to sham microglia (4.9 ± 0.8 × 10^−7^ %ID*BW). We also tested if sham injections had an impact on TSPO tracer uptake of microglia and did not find any remarkable difference in these cells when compared to control microglia of untreated age-matched mice (p = 0.999, 1-way ANOVA, **Fig. 2D**).

In the direct comparison of tumor cells and TAMs, we noted 1.37-fold higher TSPO tracer uptake of single SB28 tumor cells when compared to TAMs (p = 0.0010, paired t-test), indicating a dual cellular allocation of the tracer with even slightly higher single cell uptake of SB28 tumor cells when compared to the immune cell target population. Importantly, and highly congruent to scRadiotracing, proteomics revealed increased TSPO protein levels in SB28 tumor cells (2.98-fold, FDR-corrected p = 0.015) and TAMs (2.04-fold, p = 0.0016) when compared to control microglia. Although not reaching significance after correction for multiple comparisons, higher TSPO tracer uptake of SB28 tumor cells over TAMs was also reflected by higher TSPO protein abundance (1.46-fold, p = 0.357; **Extended Fig. 1**). We additionally proved the presence of the tracer target in SB28 tumor and immune cells by revealing that the vast majority of isolated CD11b(+) and GFP(+) cells showed TSPO co-labeling in flow cytometry (**Fig. 2E**,**F**), which is in accordance to the previously reported co-localization of TSPO and GFP/CD11b in immunohistochemistry^16^.

To determine the contributions of the distinct cell types with TSPO expression to the radioactivity in the cell pellets, we calculated a linear regression with normalized radioactivity per cell count as dependent variable and cellular proportions (GFP(+) tumor cells, CD11b(+) TAMs and ACSA2(+) astrocytes) of all enriched and depleted fractions as predictors. The regression model significantly explained the variance in single cell TSPO tracer uptake (F = 6.2, p = 0.0011, R² = 0.25, R²_adj_ = 0.21). Tumor cells showed highest contribution to the single cell TSPO tracer uptake (β = 0.76 p = 0.00017), followed by TAMs (β = 0.57, p = 0.0032) and astrocytes (β = 0.34, p = 0.0074) (**Fig. 2G**). A global model using measured radioactivity, total cell count and cell proportions indicated that 76% of the variance in radioactivity measures could be explained by the cellular component of tumor cells, TAMs and astrocytes (F = 46.6, p < 0.0001, R² = 0.772, R²_adj_ = 0.756). To question the performed calculations in depth, we correlated the radioactivity measured in the depleted tumor fractions, containing 39% ± 2% GFP(+), 7% ± 1% CD11b(+) and 16% ± 3% ACSA2(+) cells, with estimated summed up radioactivity by single cell tracer uptake and cell count (R = 0.74, p = 0.0002).

In summary, the established methodology accurately determines radiotracer uptake of single cells, which closely reflects relative protein abundance in respective cell types. The cellular allocation of TSPO as a biomarker consist of overexpression in both tumor and immune cells in SB28 tumors.

### Single cell tracer uptake explains inter-individual PET heterogeneity and reveals dominant association of tumor cells with TSPO-PET signals

To bridge the gap between cellular tracer uptake and tumor imaging, PET was added to the experimental workflow of scRadiotracing in SB28 glioblastoma mice (**Fig. 3A**). TSPO-PET indicated heterogeneity of tumor signals 2.5 weeks after surgery (range of SUV_mean_: 0.937 – 1.825), which were strongly elevated when compared to sham injection (SUV_mean_: 0.842 ± 0.032; p < 0.0001; **Fig. 3B**). We correlated the individual single cell TSPO tracer uptake of SB28 tumor cells and TAMs with the TSPO-PET signal in the lesion and observed similar degrees of strong association for both cell types (**Fig. 3C**). Interestingly, we also found an inter-correlation between single cell tracer uptake of tumor cells and TAMs (R = 0.758, p = 0.0011), which indicated bidirectional dependence of TSPO enrichment in tumor and immune cells (**Fig. 3D**). A combined vector of single cell TSPO tracer uptake of both tumor cells and TAMs strongly correlated with the magnitude of the tumor TSPO-PET signal (R = 0.848, p < 0.0001; **Fig. 3E**). A regression model of TSPO tracer uptake of tumor cells and TAMs however demonstrated that only tumor cells contributed significantly to the TSPO-PET signal (tumor cells: β = 0.55, p = 0.038; TAMs: β = 0.35, p = 0.157).

**Figure 3:**
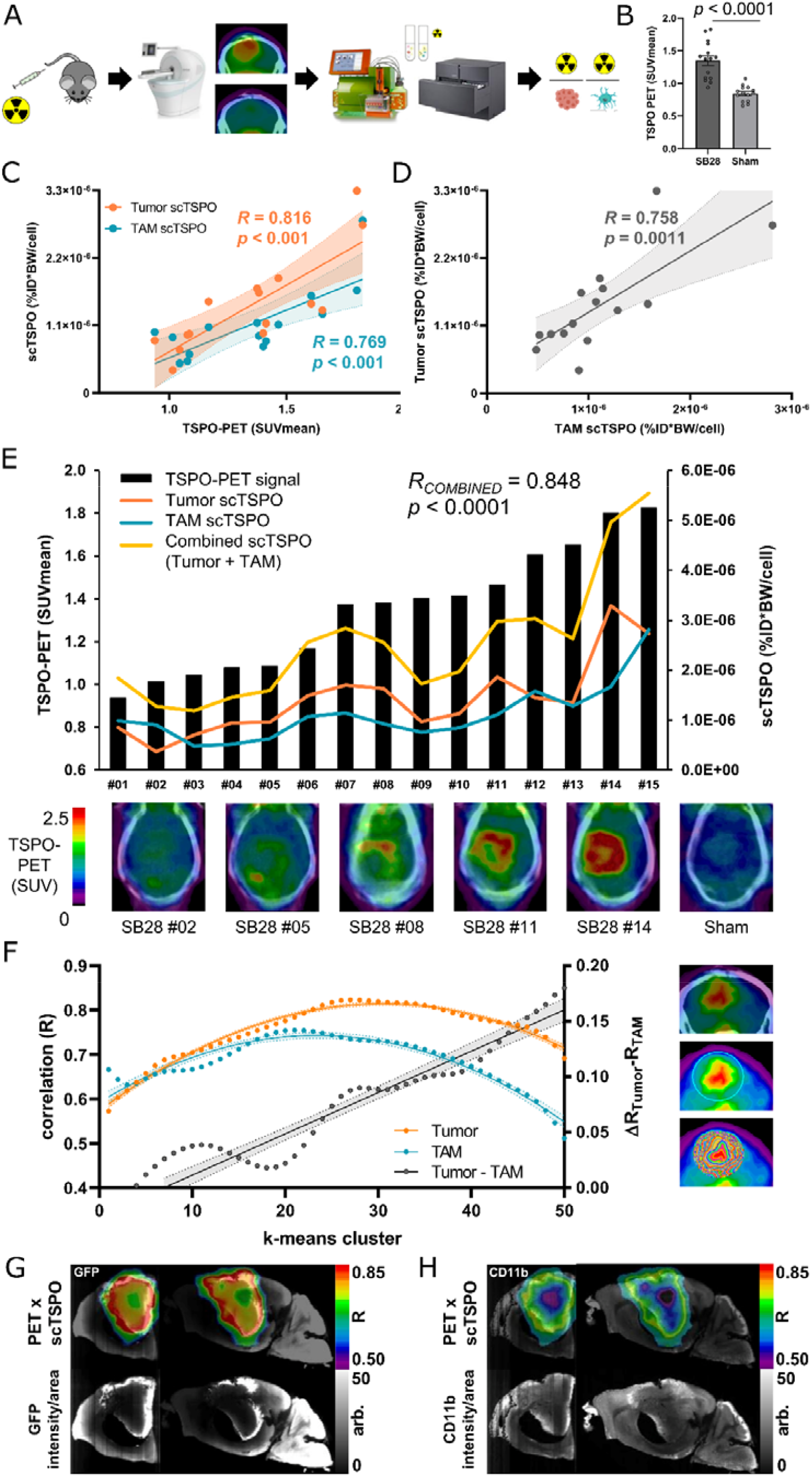
Correlation of single cell TSPO tracer uptake of tumor cells and TAMs with TSPO-PET indicates association of TSPO-PET heterogeneity with single cell tracer enrichment. **(A)** Schematic illustration of the scRadiotracing workflow in the SB28 glioblastoma mouse model including *in vivo* tracer application, PET imaging and cell sorting, resulting in radioactivity per single cell. **(B)** TSPO-PET imaging indicated significantly higher lesion site signals in SB28 tumor mice (n=15) than in sham (n=14) animals at day 18 after inoculation. Unpaired t-test, mean±SEM. **(C)** Using the established quantitative endpoint of radioactivity per single tumor cell and TAM, the correlation between scRadiotracing and TSPO-PET showed a strong dependence of PET signals from both tumor and TAM single cell TSPO tracer uptake (scTSPO, normalized to injected dose (ID) and body weight (BW)) in SB28 mice. N=15, R = Pearson’s coefficient of correlation. Error bands represent 95% confidence interval. **(D)** Inter-correlation of scTSPO of tumor cells and TAMs in the SB28 glioblastoma mouse model. N=15, R = Pearson’s coefficient of correlation. Error band represents 95% confidence interval. **(E)** Association between combined scTSPO and heterogeneous TSPO-PET signals. Black bars symbolize the individual TSPO-PET signal for all SB28 mice investigated (n=15). The curves represent scTSPO of tumor cells (orange) and TAMs (blue), both used to calculate a combined vector of cellular tracer uptake (yellow) for each individual animal. Increasing TSPO-PET signals were associated with increasing scTSPO, indicating a dependence of heterogeneous PET signals from single cell tracer uptake. Axial sections of TSPO-PET images upon the individual contrast enhanced computed tomography (ceCT) illustrate inter-individual TSPO-PET signal heterogeneity of SB28 tumors. R = Pearson’s coefficient of correlation. **(F)** Visualization of the regional k-means cluster analysis. A sphere was placed over the entire signal enhancement in TSPO-PET, which was followed by application of k-means clustering. This resulted in 50 volumes of interest (VOIs) defining 50 intratumoral regions of increasing signal intensity (images on the right). The function between increasing cluster grade and scTSPO to PET correlations of tumor cells and TAMs followed an inverted U-shape (quadratic fit) for both cell types. The correlation coefficients (R) of TSPO-PET signals with scTSPO of tumor cells showed its peak in clusters of higher rank (cluster 27; orange) when compared to the respective cluster peak of TAMs (cluster 21; blue). Increasing cluster hierarchy (i.e. the TSPO-PET hotspot) was associated with predominant dependency of TSPO-PET signal intensity from scTSPO (linear fit of ΔR_Tumor_-R_TAM_; grey). (**G**,**H**) Strong regional agreement between tumor and TAM cell density (GFP/ CD11b, light sheet microscopy) and regions with high scTSPO-to-PET associations. Coronal and sagittal slices show projections of correlation coefficients onto the k means cluster VOIs of an individual mouse. Notably, both analyses resulted in a characteristic spherical distribution of peak values that encircled the “cold” tumor core. R = Pearson’s coefficient of correlation per voxel, arb. = arbitrary units.

Going into more detail, we interrogated the regional heterogeneity of tumor TSPO-PET signals and correlated single cell TSPO enrichment with the regional PET signal magnitude. Therefore, we performed a cluster-based analysis (k-means clustering, 50 clusters defining intratumoral regions of increasing signal intensity) of TSPO-PET in the tumor lesion (**Fig. 3F**) and correlated single cell tracer uptake with the PET signal in respective clusters. The association between single cell TSPO enrichment and TSPO-PET signal intensity in the cluster followed an inverted U-shape function for tumor cells and TAMs, with higher agreement between tumor cell TSPO and regional TSPO-PET signal when compared to TAM TSPO and regional TSPO-PET signal (**Fig. 3F-H**). Strikingly, the association between TSPO-PET and single cell tracer uptake was increasingly dominated by tumor cell TSPO enrichment as a linear function of cluster hierarchy (y = 0.004x-0.026; R = 0.947; p < 0.0001; **Fig. 3F-H**). This indicates that especially regions with high TSPO-PET signal intensities were characterized by strong dependence from single tumor cell TSPO. Thus, the combination of PET and scRadiotracing identified a strong association between PET signals and cellular tracer uptake, which was characterized by a dominant dependency of TSPO-PET signals from tumor cells when compared to immune cells.

### *In vitro* scRadiotracing in human glioma underlines translational value of the methodology and confirms higher TSPO tracer uptake of tumor cells when compared to TAMs

To test for a potential translational value of single cell tracer uptake measures in human glioblastoma, we applied *in vitro* scRadiotracing in a cohort of patients with high-grade and low-grade glioma that underwent biopsy or tumor resection (**Table 1, Extended Table 1**). Tissue samples were investigated immediately after surgery applying *in vitro* [^18^F]GE-180 incubation of individual single cell suspensions and subsequent cell sorting (**Fig. 4A**). The rationale was to check for discrepancies of cellular tracer binding between species. scRadiotracing resulted in enriched fractions with visually well discernible populations of tumor cells and TAMs (**Extended Fig. 2**). High-grade glioma indicated similar heterogeneity of tumor-cell-to-TAM ratios in the single cell suspension when compared to low-grade glioma (**Fig. 4B**). As the purity of tumor and TAM enriched fractions was limited in human samples, negligibility of TSPO tracer uptake of non-tumor/non-TAM cells was proven by showing that only the number of isolated TAMs correlated with measured tracer signal in the TAM enriched cell pellet (β = 0.875, p = 0.0021), whereas non-tumor/non-TAM cells did not contribute to the overall magnitude of radioactivity in the same sample (β = 0.118, p = 0.542; **Fig. 4C**). Single cell TSPO enrichment in TAMs was similar between patients with high-grade and low-grade glioma (1.6 ± 0.9 × 10^−7^ %ID vs. 2.3 ± 1.0 × 10^−7^ %ID, p = 0.618), whereas TSPO enrichment of tumor cells was higher in patients with high-grade glioma (4.5 ± 1.3 × 10^−7^ %ID) when compared to patients with low-grade glioma (1.4 ± 0.5 × 10^−7^ %ID, p = 0.047, **Fig. 4D**). Within the group of patients with high-grade glioma, TSPO enrichment of tumor cells was stronger when compared to TAMs (p = 0.0014; **Fig. 4D**). Contrary, TSPO enrichment of tumor cells showed no significant difference when compared to TAMs within the group of patients with low-grade glioma. We questioned if previous therapy constitutes a potential confounder of single cell tracer uptake, but we did not find any significant impact of a previous therapy (radiation and/or chemotherapy, 11 out of 20) or resection (9 out of 20) on TSPO enrichment of single tumor cells or TAMs (all p > 0.05).

**Table 1:** Characteristics of the human glioma cohort. rs6971 = TSPO polymorphism rs6971^42^, LAB = low-affinity binding status, MAB = medium-affinity binding status, HAB = high-affinity binding status, Dx = Diagnosis, IDH = isocitrate dehydrogenase mutation, m = male, f = female, y = years. See **Extended Table 1** for supplemental data.

| ID | Age (y) | Sex | Dx | Disease duration (m) | WHO grade | rs6971 | IDH | Sample acquisition | Previous therapy | [<sup>18</sup>F]GE-180 TSPO-PET |
| --- | --- | --- | --- | --- | --- | --- | --- | --- | --- | --- |
| #1 | 36.0 | m | Astrocytoma | 5.6 | 2 | HAB | + | Resection | - | + |
| #2 | 59.0 | f | Oligodendroglioma | 13.6 | 2 | MAB | + | Resection | + | + |
| #3 | 50.6 | f | Glioblastoma | 0.9 | 4 | MAB | - | Resection | + | + |
| #4 | 46.9 | f | Oligodendroglioma | 0.3 | 2 | HAB | + | Resection | - | + |
| #5 | 75.7 | f | Glioblastoma | 0.2 | 4 | MAB | - | Biopsy | - | + |
| #6 | 69.1 | f | Glioblastoma | 1.0 | 4 | LAB | - | Resection | + | - |
| #7 | 64.3 | f | Oligodendroglioma | 17.8 | 2 | - | + | Biopsy | + | - |
| #8 | 59.3 | m | Astrocytoma | 2.6 | 2 | HAB | + | Biopsy | + | + |
| #9 | 36.4 | f | Oligodendroglioma | 1.1 | 2 | HAB | + | Biopsy | - | + |
| #10 | 44.0 | m | Oligodendroglioma | 12.3 | 3 | MAB | + | Biopsy | + | + |
| #11 | 72.3 | f | Glioblastoma | 0.1 | 4 | - | - | Resection | - | - |
| #12 | 39.7 | f | Astrocytoma | 4.5 | 2 | - | + | Biopsy | - | - |
| #13 | 27.2 | f | Astrocytoma | 0.7 | 2 | - | + | Biopsy | - | - |
| #14 | 68.7 | m | Glioblastoma | 0.1 | 4 | LAB | - | Resection | - | + |
| #15 | 66.8 | f | Oligodendroglioma | 25.9 | 2 | - | + | Resection | + | - |
| #16 | 58.5 | m | Astrocytoma | 20.8 | 3 | - | + | Biopsy | + | + |
| #17 | 52.8 | f | Oligodendroglioma | 20.9 | 3 | HAB | + | Biopsy | + | + |
| #18 | 49.6 | m | Oligodendroglioma | 1.2 | 3 | HAB | + | Biopsy | + | + |
| #19 | 57.9 | m | Astrocytoma | 12.9 | 3 | MAB | + | Biopsy | + | - |
| #20 | 72.0 | m | Glioblastoma | 1.4 | 4 | HAB | - | Resection | - | + |

**Figure 4:**
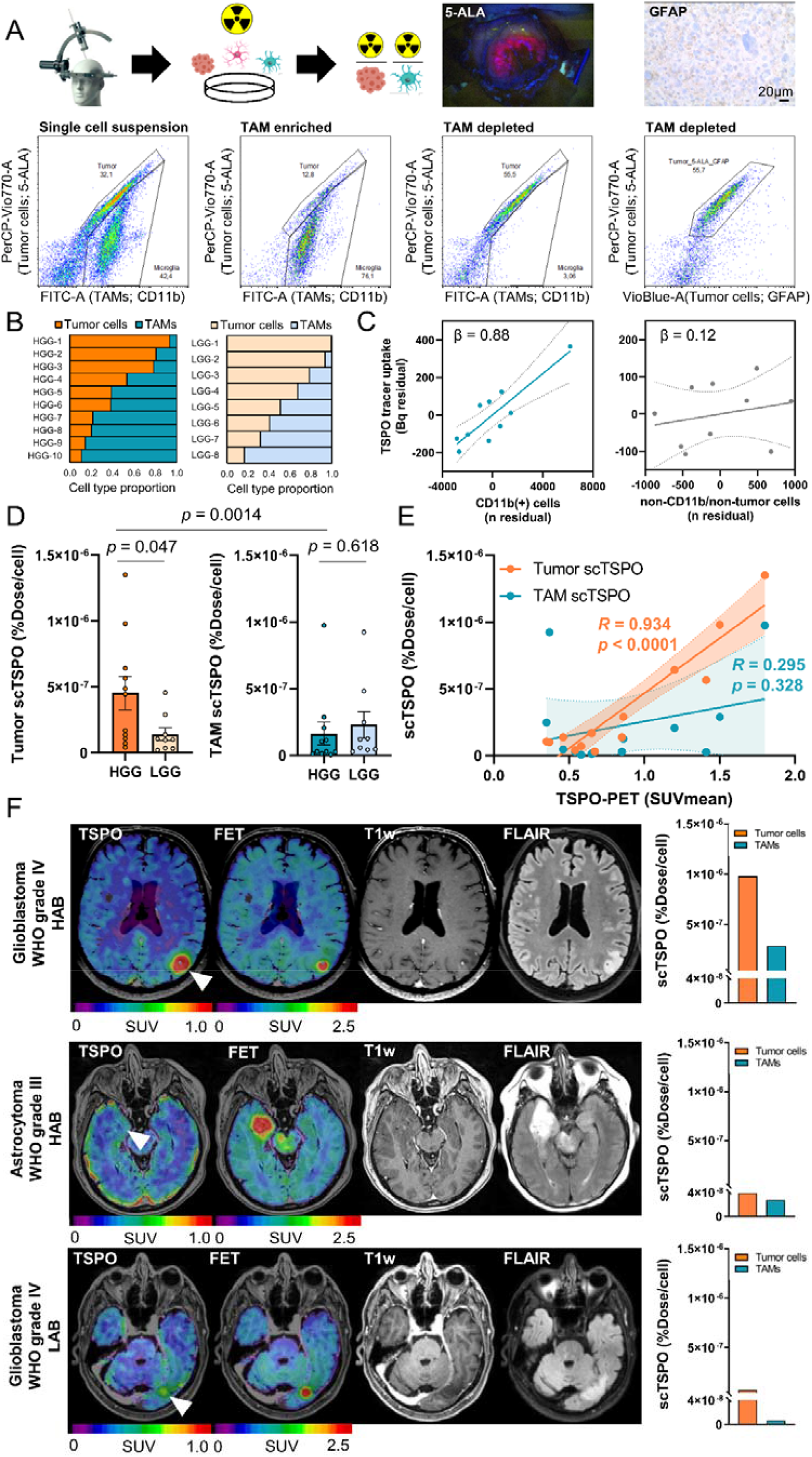
Translation of scRadiotracing to human glioma samples confirms higher TSPO tracer uptake in tumor cells and stronger contribution of tumor cells to the TSPO-PET signal when compared to TAMs. **(A)** Schematic illustration of the *in vitro* scRadiotracing workflow in human glioma samples (n=20), leading to calculation of radioactivity per single tumor cell and TAM. Pseudocolor plots derived from flow cytometry show the applied gating strategy. The single cell suspension (left) was separated into TAM-enriched (CD11b(+), second from left) and tumor enriched (third from left) fractions. Tumor cells were defined via GFAP or 5-ALA after confirmation of 5-ALA-positivity during surgery or after confirmation of GFAP-positivity during neuropathological workup of the same samples. 5-ALA positive cells co-localized with GFAP-positive cells in samples where both markers allowed tumor cell identification (right). **(B)** Relative distribution of tumor cells and TAMs in the single cell suspension of human glioma samples revealed similar heterogeneity in human high-grade glioma (HGG, n=10) compared to low-grade glioma (LGG, n=2). The initial two patients did not receive an analysis of the single cell suspension. **(C)** Regression model indicated strong contribution of TAMs (CD11b(+) cells) but not of CD11b(−)/non-tumor cells to the measured activity in the investigated TAM-enriched samples of patients that underwent biopsy (n=10). Linear regression, β = standardized regression coefficient, error bands represent 95% confidence interval. **(D)** Comparison of single cell tracer uptake (scTSPO) of tumor cells and TAMs in samples of human HGG (n=11) and LGG (n=9). Tumor cells of HGG indicated higher scTSPO than tumor cells of LGG, whereas there was no significant difference for TAMs (unpaired t-test). Tumor cells of HGG showed higher scTSPO than TAMs of HGG (paired t-test). Mean±SEM. **(E)** Correlation of TSPO-PET signals with scTSPO elucidated a strong correlation between PET signals and tumor cell TSPO enrichment and no correlation between PET signals and TAM TSPO enrichment. N=13, R = Pearson’s coefficient of correlation. Error bands represent 95% confidence interval. **(F)** Example of three patients with HGG investigated via *in vitro* scRadiotracing. All three patients showed similar signals in amino acid PET (FET-PET) and only little contrast enhancement in MRI. The patient with high tumoral TSPO-PET signal (upper row, glioblastoma, WHO grade IV, high-affinity binding status (HAB)) showed distinctly more TSPO tracer uptake in both tumor cells and TAMs when compared to the patients with only faint (middle row, astrocytoma, WHO grade III, high-affinity binding status) or low (bottom row, glioblastoma, WHO grade IV, low-affinity binding status (LAB)) tumoral signal in TSPO-PET. The patient with glioblastoma and low-affinity binding status (bottom row) showed nearly absent single cell tracer uptake of TAMs but notable TSPO tracer uptake of tumor cells. T1w = T1-weighted, FLAIR = Fluid-attenuated inversion recovery.

In accordance with the combination of PET and scRadiotracing in the SB28 mouse model, we correlated single cell TSPO enrichment with TSPO-PET imaging, which was performed in a subset of patients prior to surgery. TSPO enrichment of tumor cells (R = 0.934, p < 0.0001) but not TSPO enrichment of TAMs (R = 0.295, p = 0.328) was associated with the lesion signal in TSPO-PET (**Fig. 4E**). Individual patients with varying TSPO-PET signal intensities but similar FET and (ce)MRI findings showed a strong agreement between lesion signals and corresponding tumor cell TSPO enrichment (**Fig. 4F**). Taken together, the dominance of tumor cell radiotracer uptake and the association between scRadiotracing and PET in human tissue samples closely reflected our findings in SB28 mice. Therefore, *in vitro* scRadiotracing provides an opportunity to efficiently check for consistency in cellular target binding between murine and human samples.

### The triangle of PET, scRadiotracing and 3D-histology dissects cellular sources of regional PET signals and pinpoints dominant contribution of tumor cells to the TSPO-PET signal

Since PET tracer signals are a product of cellular tracer uptake and regional cell type abundance, we used 3D-histology via tissue clearing and light sheet microscopy (**Fig. 5A**) to determine absolute and relative cell numbers of GFP(+) tumor cells and CD11b(+) immune cells throughout entire SB28 tumors in the intact ipsilateral hemispheres. GFP(+) volumes were 5.5-fold larger when compared to CD11b(+) volumes (18.6 ± 6.8 mm³ vs. 3.4 ± 1.5 mm³, p = 0.037; **Fig. 5B**,**C**). Confocal microscopy revealed a GFP(+) volume of 4493 µm³ per SB28 tumor cell and a CD11b(+) volume of 2713 µm³ per TAM (**Fig. 5D**), resulting in higher cell count of SB28 tumor cells when compared to TAMs in individual SB28 lesions (**Fig. 5E**). Considering both single cell tracer uptake and cell distributions of tumor cells and TAMs, we calculated a tumor-cell-to-TAM contribution to the TSPO-PET signals in SB28 lesions of 3.5:1 (**Fig. 5E**).

**Figure 5:**
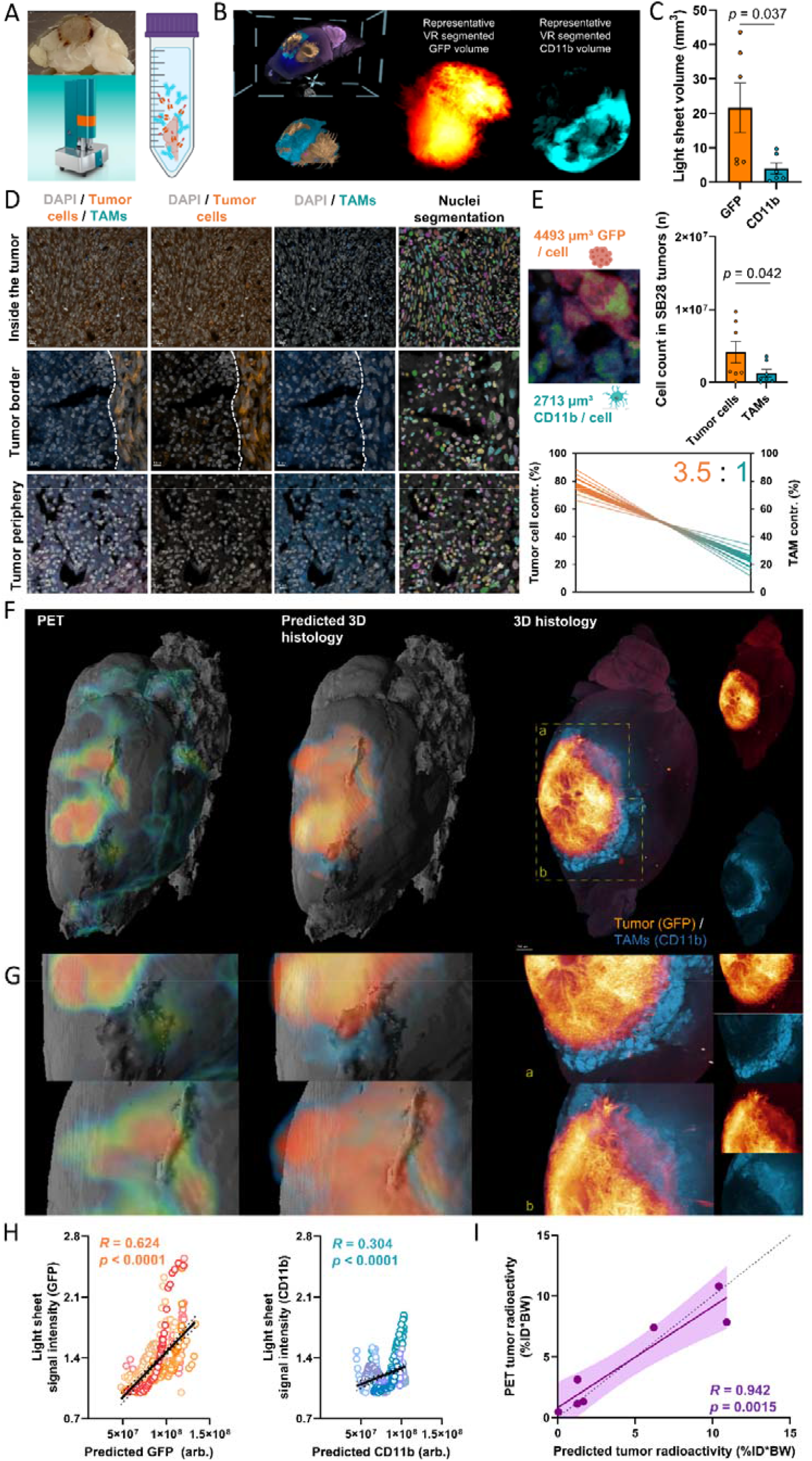
Integrated analysis of regional PET signals, cellular tracer uptake and 3D-histology via tissue clearing and light sheet imaging. **(A)** A glioblastoma inoculated brain (top left) is cleared with a modified version of 3DISCO^44^ and imaged with light sheet florescent microscope (bottom left). The details of the clearing protocol are shown on the right panel. **(B)** The imaged brains are annotated using a VR-based annotation tool, representative masks of the tumor (middle) and TAMs (right) are demonstrated. This process determined the occupied volume of GFP(+) tumor cells and CD11b(+) TAMs within the whole tumor for each individual mouse using segmentation via syGlass virtual reality tool^45^. **(C)** Quantitative comparison of determined GFP(+) and CD11b(+) volumes within individual SB28 tumors (n=7, paired t-test). **(D)** Confocal microscopy was used to determine the average GFP(+) volume per single tumor cell and the average CD11b(+) volume per single TAM by automatized count of DAPI nuclear stains within regions of exclusive GFP/CD11b expression. GFP(+) tumor cells accumulated within the tumor core (upper row), whereas the tumor periphery provided areas that were nearly exclusively composed of CD11b(+) TAMs (lower row). The right column illustrates nuclei segmentation using cellpose^46^ algorithm, which was applied to determine cell numbers per section. **(E)** Upper row: Quantitative comparison of the absolute tumor and TAM cell count after extrapolation by individual light sheet volumes (GFP-occupancy and CD11b-occupancy) and averaged GFP/CD11b volumes per single cell (left). Tumor cells revealed higher numerical abundance over TAMs in SB28 tumors (n=7, paired t-test). Lower row: Contribution of tumor cells to the overall TSPO-PET signal exceeded contribution of TAMs (3.5:1). Lines connecting tumor cell and TAM contributions indicate the matrix of variance as determined by means and SEM bands of scTSPO and volume proportions (thick lines: mean of scTSPO x upper/lower SEM Band of tumor volumes; thin lines: upper/lower SEM band of scTSPO x upper/lower SEM Band of tumor volumes). **(F)** Regional TSPO-PET signals (left) were combined with single cell tracer uptake values of tumor cells and TAMs to predict cell type abundance within individual SB28 tumors. Predicted 3D-histology (middle) shows high regional agreement with standard of truth 3D-histology as obtained by light sheet microscopy (right). The whole set of analyzed mice is presented in **Extended Fig. 3.** **(G)** Representative magnified areas within the SB28 tumor underline the regional agreement between predicted and standard of truth 3D histology. **(H)** Quantitative regional correlation of predicted and standard of truth 3D-histology within k means clusters (**Fig. 3**) shows high agreement of predicted tumor cell abundance and moderate agreement of predicted TAM abundance. Light sheet TIFFs were spatially co-registered to PET as 3D RGB images to extract signal intensities of light sheet as a surrogate of cell type abundance in all PET derived clusters. To predict 3D histology (separately for GFP and CD11b), PET signal intensities in each cluster were multiplied with the cluster-based PET correlation coefficient (**Fig. 3F**) and divided by single cell tracer uptake. n=350 single regions from n=7 tumors. R = Pearson’s coefficient of correlation. Error bands represent 95% confidence interval. arb. = arbitrary units. **(I)** PET measured radioactivity of the tumor was predicted by the radioactivity derived from the product of cell type abundance (tumor cells and TAMs) and single cell tracer uptake in all individual mice that underwent PET imaging and light sheet microscopy. R = Pearson’s coefficient of correlation. Error bands represent 95% confidence interval. Dashed line represents line of identity (y=x).

Next, PET, scRadiotracing and 3D-histology were used as an integrated concept. Importantly, the regional resolution of 3D-histology could be predicted by combining lesion PET signals with single cell tracer values (**Fig. 5F**,**G, Extended Fig. 3**). For this purpose, we used the regional association between single cell tracer uptake of SB28 tumor cells and TAMs with TSPO-PET (i.e. correlation coefficients per cluster, **Fig. 3F**) as a cluster-based PET weighting factor. k-means cluster PET signals of all mice that received TSPO-PET imaging and 3D-histology were multiplied with their respective correlation coefficients and divided by single cell tracer uptake of both tumor cells and TAMs (**Fig. 2D**) to predict individual regional abundance (i.e. light sheet signal intensity) for each cell type (**Fig. 5F-H**). Predicted and standard of truth 3D-histology were strongly correlated (tumor cells: R = 0.624, p < 0.0001; TAMs: R = 0.304, p < 0.0001; **Fig. 5H; Extended Fig. 3**). Hot spots of GFP-positivity were co-localized with regions of high TSPO-PET signal intensity (**Fig. 5G**), fitting to the strong association between tumor cell TSPO and regional TSPO-PET (**Fig. 3F**). The radioactivity in individual tumors as measured by PET was precisely predicted by aggregated single cell tracer uptake and cell count of tumor cells and TAMs (**Fig. 5I**). In summary, exploiting the established methodological combination of PET, scRadiotracing and 3D-histology enables to pinpoint complex PET signals to their detailed cellular origins.

### Proteomics identifies potential TAM-specific radiotracer targets with superior binding capacity when compared to TSPO

Since tumor cells dominate the TSPO-PET signal, the development of novel PET radiotracers with higher specificity for TAMs could provide added value to glioblastoma diagnostics. We therefore generated murine proteome data to identify potential TAM-specific radiotracer targets. Starting from a total of 7869 proteins relatively quantified between TAMs and microglia of healthy control mice, 1097 showed higher fold-changes than TSPO (**Fig. 6A**,**B**). 165 of these proteins could not be quantified in SB28 tumor cells and additional 16 proteins showed >10-fold higher levels in TAMs when compared to SB28 tumor cells (**Fig. 6B**), thus providing sufficient specificity for TAM over tumor cell expression.

**Figure 6:**
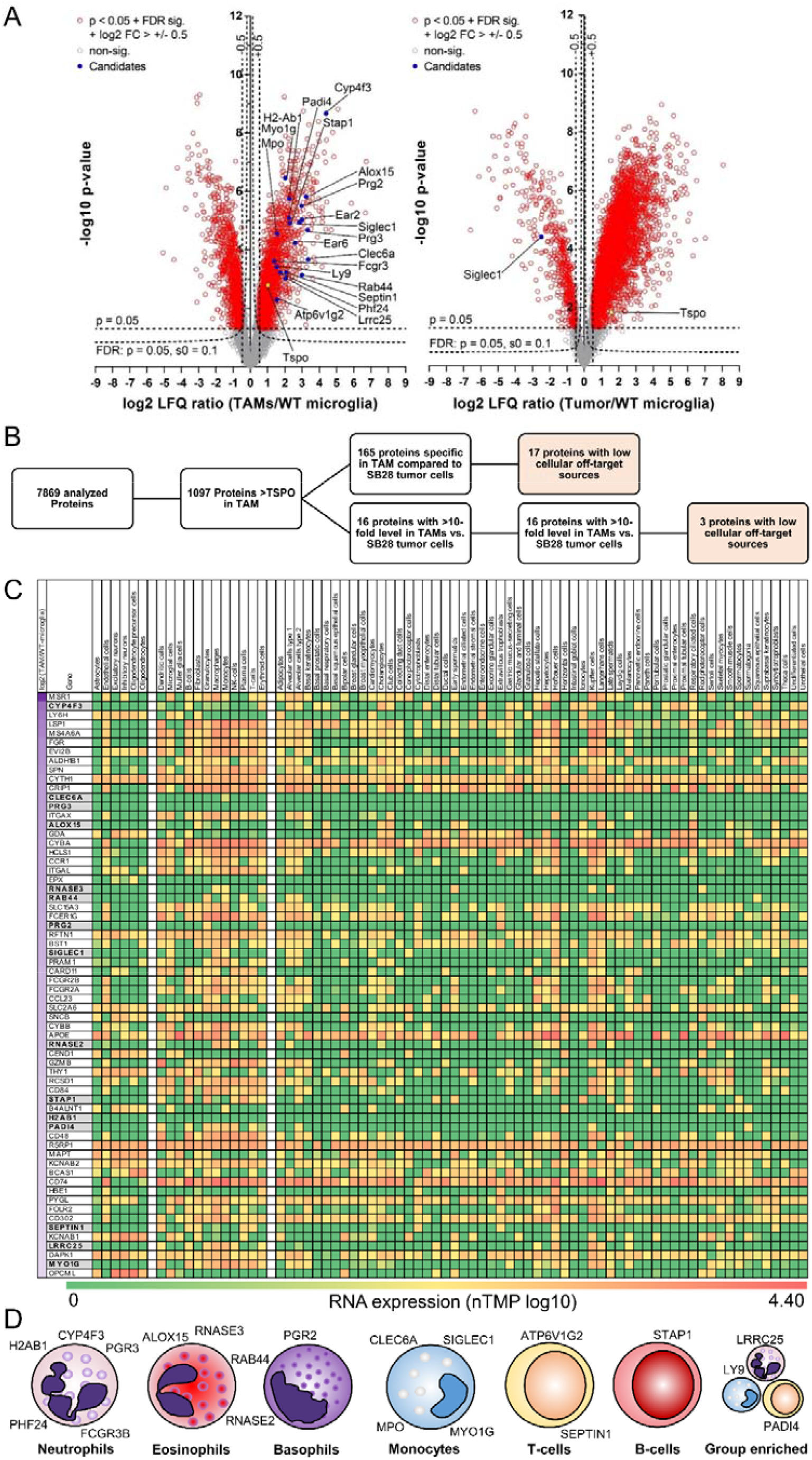
Proteome analysis of the SB28 tumor microenvironment identifies TAM-specific radiotracer targets for glioblastoma. **(A)** Volcano plot representation of the different protein levels in isolated TAMs (n = 5) and isolated SB28 tumors cells (n = 3) in comparison with control microglia isolated from age matched mice (wt-microglia; *n* = 6). TSPO (yellow) showed intermediate elevation of protein levels in TAMs and SB28 tumor cells. 1098 out of 7869 analyzed proteins indicated higher protein levels in TAMs than did TSPO. **(B)** Selection process of potential radiotracer targets for specific detection of TAMs over SB28 tumor cells. 165 of the identified proteins in TAMs were not detected in SB28 tumor cells and another 16 proteins had >10-fold higher levels in TAMs compared to SB28 tumor cells. A total of 20 proteins showed low cellular off-target sources and were identified as potential TAM radiotracer targets. **(C)** Heat-map screening for TAM-specificity of the 181 proteins of interest. The Human Protein Atlas^47^ was used to determine RNA expression levels of all proteins of interest in i) resident off-target cells of the brain (left cell type columns), ii) resident and infiltrating cells in the presence of glioblastoma (middle cell type columns) and iii) off-target cells of the organism (right cell type columns). Genes are sorted by protein level differences in TAMs compared to control microglia of untreated mice (top to bottom; log2 label-free quantification ratio; first column). 64 proteins with highest elevation (log2 (TAMs/wt-microglia) > 2, left column) are illustrated in **C** and all proteins are provided in **Extended Fig. 4**. Proteins of interest are highlighted in grey. **(D)** Immune cell expression cluster allocation of proteins of interest. scRNA data of the Human Protein Atlas was used to categorize the final set of identified proteins.

To translate these findings from mouse to human, we used the Human Protein Atlas^17^ to identify expression of respective human gene homologs in off-target cells in brain as an index of TAM-specificity (see first set of columns in **Fig. 6C, Extended Fig. 4**). Physiological expression in resident and invading cells of the tumor microenvironment (see second set of columns in **Fig. 6C, Extended Fig. 4**) was characterized. In a next step, we ensured low target expression in other cells in the organism to prevent peripheral radiotracer absorbance, since this may cause a sink effect and high radiation exposure (see third set of columns in **Fig. 6C, Extended Fig. 4**). This selection process resulted in 20 proteins with highly suitable characteristics as specific immune PET tracer targets (**Fig. 6D**), which were ultimately categorized into their immune cell expression cluster (**Fig. 6D**). *CYP4F3, H2AB1, PGR3, FCGR3B* and *PHF24* were allocated to the neutrophil expression cluster. *ALOX15 and RNASE3* were associated with the eosinophil expression cluster and *PGR2* was localized in the basophil expression cluster, while both expression clusters were shared for *RAB44* and *RNASE2*. An association with the monocyte expression cluster was identified for *CLEC6A, MPO, MYO1G* and *SIGLEC1*. Few identified genes belonged to B-cell (*ATP6V1G2* and *SEPTIN1*) and T-cell (*STAP1*) expression clusters and group enrichment was found for *LRRC25, LY9* and *PADI4* (**Fig. 6D**). High protein abundance, which allows sufficient PET signal-to-noise-ratios, was observed for *ALOX15, PRG2, SIGLEC1, MYO1G* and *MPO* (Label-free quantification (LFQ) intensity > 18, **Extended Fig. 5**). Radiolabeled ligands with binding affinity to these proteins may re-enter the workflow of scRadiotracing, 3D-histology and human translation to test for their cellular specificity *in vivo*.

## Discussion

We present the first study that determines *in vivo* radiotracer uptake in the complex TME of glioblastoma at cellular resolution. After validation of scRadiotracing for the immune target TSPO, we deciphered the cellular sources of TSPO-PET signals in experimental glioblastoma and patients with glioma. The established methodology accurately determined higher radiotracer uptake of single glioblastoma tumor cells when compared to immune cells, which closely reflected protein levels of respective cell types. The combination of PET and single cell tracer uptake served to predict 3D-histology. Ultimately, a detailed proteome analysis facilitated identification of TAM-specific radiotracer targets beyond TSPO.

As a prerequisite of the main analysis, a methodological validation of TSPO radiotracer measures in single cells had to be addressed. We applied a microglia depletion paradigm^18^ to compare the signal reduction observed by *in vivo* [^18^F]GE-180 TSPO-PET of mice after CSF1R inhibition with the total radioactivity attributable to the microglia cell population after scRadiotracing. Here, we confirmed that scRadiotracing was able to closely recover the tracer signal reduction observed in PET (**Fig. 1**). By CSF1R inhibition with PLX5622, microglia cells can be depleted up to 99%, which may be maintained under continuous treatment for up to 2 months^19^, but CSF1R inhibition does not change the TSPO gene expression of other CNS cells^20^. Thus, we claim that the reduction of TSPO-PET derived from the nearly complete depletion of microglia after 7-weeks PLX5622 treatment. Furthermore, we observed 12-fold higher single cell TSPO tracer uptake of microglia when compared to astrocytes, pinpointing specificity of [^18^F]GE-180 to microglia in untreated mice.

Additionally, the high congruence between tracer signal in the microglia population and PET signal reductions after microglia depletion fits to previous investigations that outlined a high correlation between [^18^F]GE-180 TSPO-PET and the microglia markers Iba1 and CD68 in immunohistochemistry^21^.

We were able to further validate scRadiotracing in tumor cells and TAMs of the experimental SB28 glioblastoma model by quantification of relative cellular protein levels of TSPO in respective cell populations using proteomics. The application of proteomics was particularly important since transcriptomics does not necessarily reflect the amount of protein that can be targeted by the radiotracer^22^. We also found that the inter-individual heterogeneity of SB28 and human tumor PET signals was correlated with the magnitude of tracer uptake per each single cell, which implies that the PET signal is mainly driven by cellular uptake and not by confounding alterations of the blood-brain-barrier^23^ or off target binding^24^. Our observations of heterogeneous single cell tracer uptake fit to the biological variability of TSPO which has not only been outlined in glioblastoma^7,25^ but also in neurodegenerative diseases^26,27^. Ultimately, the combination of PET and scRadiotracing with 3D-histology confirmed the methodological capability to quantify cellular sources of *in vivo* PET signals. In this regard, PET tracer signals are not only a function of cellular uptake but a product of cell type abundance and single cell tracer uptake, making the novel combination of scRadiotracing and 3-dimensional assessment of specific cell counts^28^ a major achievement of the current work. Furthermore, scRadiotracing allowed to predict 3D-histology at the individual level, which may have clinical impact in patients with glioma, where histology can only be acquired in subsamples of the highly heterogeneous tumor mass^29,30^.

In this study, we used TSPO as an established immune PET biomarker in neurological and neuro-oncological diseases with controversially discussed cellular signal sources. Although several correlative investigations questioned the cellular sources of TSPO tracer signals in glioma^6,31^, direct measures of radiotracer allocation have not yet been performed and contributions of tumor and immune cells to the PET signal remained unclear. Our data indicated higher enrichment of [^18^F]GE-180 in SB28 tumor cells when compared to TAMs after *in vivo* tracer administration and similar results were obtained in human high-grade glioma cells when compared to TAMs of the same tumor after *in vitro* tracer incubation. These findings were supported by stronger agreement of tumor TSPO-PET signals with individual tumor cell tracer uptake in mice and humans. 3D-histology in SB28 mice moreover showed higher abundance of tumor cells when compared to TAMs, pinpointing the dominance of tumor cells to SB28 tumor TSPO-PET signals. Still, potential discrepancies between murine and human glioblastoma (e.g. inoculation versus spontaneous tumor growth) need to be considered. Although the SB28 model is reported to closely mimic the human TME^15^ and our translational data indicate similar results regarding single cell TSPO enrichment across species, the small sample size of the human analysis in respect of large heterogeneity in diagnoses and previous therapy strategies remains to be acknowledged. Further investigations in pre-defined subgroups with larger sample sizes may aid to disentangle and differentiate therapy-related effects on TSPO expression in cells of the human TME, i.e. after radiation therapy^32^.

A high contribution of tumor cells to the TSPO-PET signal of glioblastoma is in agreement with preclinical investigations that used TSPO-PET and autoradiography after implantation of GL261 cells into a full TSPO knock-out ^31,33^. However, several preclinical studies have revealed a distinct co-localization of TSPO and Iba1/CD68 in immunohistochemistry in different glioblastoma mouse models and concluded that TAMs contribute significantly to the TSPO-PET signal^7-10^. While some of these studies have already drawn a qualitative comparison between immunohistochemistry and TSPO-PET quantification by various ligands^8-10^, our study provides first direct evidence for simultaneous allocation of a TSPO-PET tracer in both cell types. Future research should also compare single cell TSPO enrichment across the broad spectrum of different glioblastoma models (i.e. GL261) and primary human glioblastoma cell lines.

Various signal sources with a significant impact on the TSPO-PET signal in the TME call for detailed interrogation of TSPO as a glioblastoma imaging biomarker. First, lacking specificity of TSPO-PET signals for TAMs in glioblastoma hampers the biomarker’s application as a mere immune index. However, since TSPO-PET signals allow prognostication of overall survival^34^, TSPO-PET still adds valuable biomarker features to the toolbox of glioblastoma imaging assessments^35^. These quantitative TSPO-PET data even exceeded the prognostic value of gene expression levels^36^, which could be related to the direct quantification of TSPO receptor abundance instead of indirect mRNA measures. The pilot dataset of *in vitro* scRadiotracing in patients with glioma already showed feasibility even with small amounts of tissue obtained from biopsy. Thus, the current results should encourage to investigate the separate prognostic value of single cell TSPO enrichment in tumor and immune cells since such data would potentially result in different treatment strategies. Importantly, distinct binding of radioligands to different cells of the TME will also impact personalized medicine, including direct targeting of various tumors with radiotheranostics.

Beyond TSPO, TAM-specific radioligands for assessment of immune cells in the TME could facilitate monitoring of novel immunotherapies^37^. We used our murine proteomics dataset to classify TSPO within the spectrum of potential radiotracer targets. Since TSPO was only moderately elevated in TAMs and surpassed by the TSPO expression in tumor cells, there is a major need for further improvement, identification and development of TAM-specific radiotracers. The identified targets need to be further evaluated in terms of ligand accessibility and stable target expression and screened for available crystal structures or known binders that aid generation of lead molecules for radiotracer development. Importantly, several of the identified proteins matched with signature genes of immunosuppressive (i.e. *ALOX15*^38^) TAM phenotypes or predicted therapy response upon checkpoint inhibition (i.e. *LY9*^39^ or *MPO*^40^), thus indicating their potential as biomarkers of immunomodulation in cancer. scRadiotracing will provide the opportunity for a precise determination of the *in vivo* specificity of potential radioligands for these targets in their developmental process despite the complexity of the TME.

In a nutshell, the established methodology yields robust and reliable quantification of cellular radiotracer uptake in mice and patients and will serve to disentangle cellular sources of PET signals of established or newly developed tracers. This is of particular importance since many potential radiotracer targets are shared between tumor cells, TME cells and resident cells of the brain^41^.

## Methods

### Study design

The primary goal of the study is to resolve PET tracer signals at cellular resolution. We challenged our novel methodology by targeting TSPO as a biomarker with complex cellular sources in glioblastoma. For this purpose, a combination of radiotracer injection and immunomagnetic cell sorting (MACS) was applied (scRadiotracing^11^). To validate the workflow, the MACS data of microglia in untreated mice were compared to microglia depletion experiments using CSF1R inhibition in combination with [^18^F]GE-180 TSPO-PET. In mice with orthotopic implanted tumor (SB28, 2.5 weeks post inoculation) or sham injection, the brain was removed 75 minutes after [^18^F]GE-180 injection and whole-brain activity was measured. The tumor or the injection site was dissected, followed by radioactivity measurement and dissociation of the tumor tissue into a single cell suspension. Tumor cells (negative selection) and TAMs (CD11b) were isolated via magnetic cell sorting using specific antibodies. The radioactivity in the cell-specific pellets was measured by a highly sensitive gamma counter. To determine the absolute cell number in each pellet as well as the proportions (i.e. purity) of the different cell populations, flow cytometry was performed subsequent to cell sorting. TSPO tracer uptake was determined per single cell and compared to TSPO protein expression levels, whereas contributions of the different cell populations to the total activity were calculated by multiple regression. The methodology was translated to human resection and biopsy samples of patients with high-grade and low-grade glioma and *in vitro* scRadiotracing was performed to determine proportions of tumor and immune cell tracer uptake. To acknowledge PET signals as product of cellular uptake and cellular abundance, 3D-histology via light sheet microscopy was performed in PFA perfused mouse brains to obtain absolute cell counts and relative quantitative proportions of glioblastoma and immune cells within the tumor. Then, an integrated model of PET, single cell tracer uptake and 3D-histology was used to disentangle regional PET signals by their distinct cellular components. Finally, TSPO protein levels were characterized within the whole spectrum of the tumor and immune cell proteome, with the goal to determine TAM-specific radiotracer targets.

### Animals

All animal experiments were performed in compliance with the National Guidelines for Animal Protection, Germany and with the approval of the regional animal committee (Government of Upper Bavaria) and overseen by a veterinarian. All animals were housed in a temperature- and humidity-controlled environment with a 12-h light–dark cycle, with free access to food (Ssniff, Soest, Germany) and water.

For scRadiotracing experiments, eight-week-old C57BL/6 mice were purchased from Charles River (Sulzfeld, Germany) and acclimated for at least 1 week. At day 0, the mice were inoculated with 100,000 SB28-GFP cells suspended in 2 µl of DMEM (Merck, Darmstadt, Germany) (glioblastoma mice, n=27) or 2 µl of saline (sham mice, n=14). Additional n=8 mice received no treatment before scRadiotracing, serving as control animals. For inoculation, mice were anesthetized with intraperitoneal (i.p.) injections of 100 mg/kg ketamine 10% and 10 mg/kg xylazine 2% in 0.9% NaCl. Anesthetized mice were immobilized and mounted onto a stereotactic head holder (David Kopf Instruments, Tujunga, CA, USA) in the flat-skull position. After surface disinfection, the skin of the skull was dissected with a scalpel blade. The scull was carefully drilled with a micromotor high-speed drill (Stoelting Co, Wood Dale, IL, USA) 2 mm posterior and 1 mm left of the bregma. By stereotactic injection, 1 × 10^5^ cells were applied with a 10 µl Hamilton syringe (Hamilton, Bonaduz, Switzerland) at a depth of 2 mm below the drill hole. Cells were slowly injected within 1 minute and after a settling period of another 2 minutes the needle was removed in 1 mm steps per minute. After that, the wound was closed by suturing. Mice were checked daily for tumor-related symptoms and sacrificed when tumor burden (i.e. appearance, coordinative deficits, motor symptoms) reached stop criteria (not reached in any animal). On the last day of the experiment, the mice were injected with 15 ± 1 MBq [^18^F]GE-180 into the tail vein before cervical dislocation and brain extraction at 75 minutes post injection (n=33). N=15 of these glioblastoma and all sham mice received TSPO-PET imaging directly before brain extraction. Additional n=7 glioblastoma mice received TSPO-PET imaging and perfusion with 4% PFA prior to 3D-histology by light sheet microscopy. Another n=14 C57BL/6 mice were used for extraction of naïve control microglia (n=8 scRadiotracing; n=6 proteomics).

For validation of scRadiotracing as a reliable method to determine TSPO-PET signal sources at cellular resolution, we analyzed combined data of pharmacological depletion of microglia by CSF1R inhibition^48^ which was performed in C57BL/6 mice of two datasets^13,14^ by PLX5622 (1200 ppm, n=15) and vehicle controls (n=14), aged 10.1 ± 2.1 months. The TSPO-PET scan was performed in the last week of seven weeks treatment.

### Cell culture

SB28-GFP cells^49^ were cultured in DMEM containing MEM non-essential amino acids (1x), 1% Penicillin-Streptomycin solution (Thermo Fisher Scientific, Waltham, MA, USA) and 10% fetal bovine serum (FBS, Biochrome, Berlin, Germany). Cell cultures were maintained in the incubator at 37°C in humidified and 5% CO_2_-conditioned atmosphere. Cells were passaged when the cell density in the flask reached 80% confluence.

### Radiosynthesis

Automated production of [^18^F]GE-180 was performed on a FASTlab™ synthesizer with single-use disposable cassettes. The pre-filled precursor vial was assembled on the cassette and the cassette was mounted on the synthesizer according to the set-up instructions. The FASTlab™ control software prompts were followed to run the cassette test and to start the synthesis. No-carrier-added ^18^F-fluoride was produced via ^18^O(p, n)^18^F reaction by proton irradiation of ^18^O-enriched water and delivered to the ^18^F incoming reservoir. The fully automated manufacturing process consists of the following steps: trapping of ^18^F-fluoride on a QMA cartridge, elution using Kryptofix^®^222, potassium hydrogen carbonate, water and acetonitrile, azeotropic drying of ^18^F-fluoride at 120°C for 9 minutes, labelling of the precursor in MeCN at 100°C for 6 minutes, dilution of the crude product with water, tC18 cartridge based purification by use of 20 mL 40% (v/v) Ethanol and 11.5 mL 35% (v/v) Ethanol, elution of the product with 3.5 mL 55% (v/v) Ethanol and final formulation with phosphate buffer. RCY 39±7% (n=16) non d. c., synthesis time 43 minutes, RCP ≥98%.

### Immunomagnetic cell separation

MACS was performed as described previously^13^ with slight modifications for additional tumor cell isolation. Detailed descriptions of brain dissociation and isolation of different cell types were as follows.

### Mouse brain dissociation

Adult mouse brains were removed after cervical dislocation at 75 minutes post injection (p.i.) and stored in cold D-PBS. The brains were cut into small pieces and dissociated using gentleMACS Octo Dissociator with Heaters (Miltenyi Biotec, 130-096-427) in combination with different Dissociation Kits according to the manufacturer’s instructions. Adult Brain Dissociation Kit, mouse and rat (Miltenyi Biotec, 130-107-677) was used for adult mouse brain dissociation of WT mice and contralateral hemispheres of glioblastoma mice. Tumor Dissociation Kit (Miltenyi Biotec, 130-096-730) was used for dissociation of tumor tissue. The dissociated cell suspension was applied to pre-wet 70-μm Cell Strainer (Miltenyi Biotec, 130-110-916). The cell pellet was resuspended using cold D-PBS and cold debris removal solution. Cold D-PBS was gently overlaid on the cell suspension and centrifuged at 4°C and 3000g for 10 minutes with acceleration at 9 and deceleration at 5. The two top phases were removed entirely. The cell pellets were collected. Non-tumoral cell pellets were additionally resuspended with 1 mL of cold red blood cell removal solution followed by 10 minutes incubation. Cell pellets were collected for further applications.

### Isolation of tumor cells

Tumor Cell Isolation Kit, mouse (Miltenyi Biotec, 130-110-187) was used according to the manufacturer’s instructions. The prepared cell pellets were resuspended in 80 µl of D-PBS– 0.5% bovine serum albumin (BSA) buffer per 10^7^ total cells including red blood cells. 20 µl of Non-Tumor Cell Depletion Cocktail were added and the suspension was incubated at 4°C in the dark for 15 minutes. The volume was adjusted to 500 µl per total 10^7^ cells with D-PBS– 0.5% BSA buffer before proceeding to magnetic separation. The pre-wet LS columns (Miltenyi Biotec, 130-042-401) were placed at QuadroMACS Separator (Miltenyi Biotec, 130-090-976). The cell suspensions were applied onto the column. The columns were washed with 2 × 1 mL of D-PBS–0.5% BSA buffer. The flow-through containing the unlabeled cells was collected as the tumor-cell-enriched fractions. The columns were removed from the magnetic field, and the non-tumor cells were flashed out using 3 mL of D-PBS–0.5% BSA buffer.

### Isolation of TAMs and microglia

TAMs (glioblastoma mice) or microglia (sham injected or untreated control mice) were isolated from animals using CD11b MicroBeads, human and mouse (Miltenyi Biotec, 130-049-601) and a MACS separation system (Miltenyi Biotec) as described previously^50,51^. For murine samples, the prepared cell pellets were resuspended in 90 µl of D-PBS–0.5% BSA buffer per 10^7^ total cells. Ten microliters of CD11b MicroBeads per 10^7^ total cells were added and incubated for 15 minutes in the dark at 4°C. Human samples were resuspended in 80 µl of D-PBS–0.5% BSA buffer per 10^7^ total cells and 20 µl of CD11b MicroBeads per 10^7^ total cells were added and cells were incubated for 15 minutes in the dark at 4°C. Cells were washed by adding 1 to 2 mL of buffer per 10^7^ cells and centrifuged at 300g for 10 minutes. The cell pellets were resuspended in 500 µl of D-PBS–0.5% BSA. The pre-wet LS columns (Miltenyi Biotec, 130-042-401) were placed onto a QuadroMACS Separator (Miltenyi Biotec, 130-090-976). The cell suspensions were applied onto the column. The columns were washed with 3 × 3 mL of D-PBS–0.5% BSA buffer. The flow-through containing the unlabeled cells was collected as the microglia-depleted fractions. The columns were removed from the magnetic field, and microglia were flashed out using 5 mL of D-PBS–0.5% BSA buffer.

### Isolation of astrocytes

Adult Brain Dissociation Kit, mouse and rat (Miltenyi Biotec, 130-107-677) was used according to the manufacturer’s instructions. The prepared cell pellets were resuspended in 80 µl of AstroMACS separation buffer (Miltenyi Biotec, 130-117-336) per 10^7^ total cells. 10 μl of FcR blocking reagent were added and incubated for 10 minutes in the dark at 4°C. 10 μl of Anti-ACSA2 MicroBeads were added and incubated for 15 minutes in the dark at 4°C. Cells were washed by adding 1 mL of AstroMACS separation buffer and centrifuged at 300g for 5 minutes. Cell pellets were resuspended in 500 μl of AstroMACS separation buffer. The pre-wet MS columns (Miltenyi Biotec, 130-042-201) were placed at OctoMACS Separator (Miltenyi Biotec, 130-042-109). The cell suspensions were applied onto the column, followed by washing with 3 × 500 µl of AstroMACS separation buffer. The flow-through was collected containing non-astrocytic cells as an astrocyte-depleted fraction. The columns were removed from the magnetic field, and the astrocytes were flashed out using 3 mL of AstroMACS separation buffer.

### Gamma emission measurements

Radioactivity concentrations of cell pellets were measured in a gamma counter (Hidex AMG Automatic Gamma Counter, Mainz Germany), cross-calibrated to the activity in the whole brain, with decay correction to time of tracer injection for final activity calculations.

### Flow cytometry

Flow cytometry staining was performed at 4 °C. After gamma emission measurement, the cell suspension was centrifuged at 400g for 5 minutes and the supernatant was aspirated completely. The cell pellet was then resuspended in 100 µl of cold D-PBS containing fluorochrome-conjugated antibodies recognizing mouse CD11b and ACSA2 (Miltenyi Biotec, 130-113-810 and 130-116-247) in a 1:100 dilution and incubated for 10 minutes at 4°C in the dark. Samples were washed with 2 mL of D-PBS and centrifuged for 5 minutes at 400g.

Finally, cell pellets were resuspended in 500 μl of D-PBS and samples were immediately used for flow cytometry using a MACSQuant® Analyzer. Precision Count Beads (Biolegend, 424902) were added for counting the absolute number of cells for the samples of n=9 mice measured with a BD LSR Fortessa Cell Analyzer (BD Biosciences, Franklin Lakes, NJ, USA). Acquired data included absolute cell numbers and purity of GFP(+), CD11b(+) and ACSA2(+) cells in each sample.

TSPO co-staining was performed for tumor and CD11b enriched cell fractions of n=3 SB28 mice. To this end, cells were permeabilized after initial flow cytometry using Inside Stain Kit (Miltenyi Biotec, 130-090-477) according to the manufacturer’s instructions. Cell pellets were resuspended in 250 µl of cold D-PBS, 250 µl of Inside Fix Solution was added, cells were incubated for 20 minutes in the dark at 4°C, then centrifuged at 300g for 5 minutes. Complete aspiration of supernatant was followed by washing with 1 mL of cold D-PBS and another centrifugation step at 300g for 5 minutes. Cell pellets were washed with 1 mL of Inside Perm Solution and centrifuged at 300g for 5 minutes. Supernatant was aspirated, cells were resuspended in 47 µl of Inside Perm Solution and stained with 3 µl of Anti-PBR antibody [EPR5384] (abcam, ab199836). After 10 minutes of incubation, cells were washed by adding 1 mL of Inside Perm Solution, centrifuged at 300g for 5 minutes and resuspended in 300 µl cold D-PBS for flow cytometry analysis.

For subsequent proteome analysis (see below), cell pellets of n=3 tumor enriched and n=5 CD11b enriched fractions of SB28 tumor samples were stored at -80 degree together with n=6 CD11b enriched fractions (i.e. control microglia) derived from an independent cohort of age- and sex matched untreated control mice.

### Calculation of single cell TSPO tracer signal

Measured radioactivity (Bq) of cell pellets was divided by the specific cell number in the pellet resulting in calculated radioactivity per cell. Radioactivity per cell was normalized by injected radioactivity and body weight (i.e. %ID*BW). Published cell numbers of microglia^12^ were used to extrapolate the whole brain radioactivity located in the microglia fraction of untreated mice.

For tumor probes, cell numbers (tumor cells and TAMs) determined by light sheet microscopy were multiplied with single cell %ID*BW as an estimate of cell fraction specific contributions to the PET signal.

### Human samples and in vitro scRadiotracing

Human tumor tissue samples were acquired during neurosurgical biopsy or open resection and stored in Tissue Storage Solution (Miltenyi Biotec, 130-100-008) for 2-30 hours until further processing for scRadiotracing. Details on the patient cohort are provided in the results section (**Table 1**). Tissue was manually cut into smaller pieces if necessary and dissociated using Tumor Dissociation Kit, human (Miltenyi Biotec, 130-095-929). Removal of debris and red blood cells was performed for tumor resections as described above. Single cell suspensions were incubated *in vitro* at a concentration of 56 ± 8 MBq [^18^F]GE-180 in 1 mL for 30 minutes, then washed twice with 3 mL cold D-PBS and centrifuged at 400g for 5 minutes. Cell pellets were resuspended in 100 µl cold D-PBS and stained with fluorochrome-conjugated antibodies recognizing human CD11b and GFAP (Miltenyi Biotec, 130-113-810 and 130-123-846) as described above. Gamma counter measurement and flow cytometry analysis of the stained single cell suspension were performed using 10% of the probe after resuspension in 500 µl. Subsequently, TAM were isolated using CD11b microbeads (Miltenyi Biotec, 130-093-634), followed by further gamma counter measurement and flow cytometry. Proportions and cell count of TAM in the resulting cell pellets were determined using CD11b. For tumor cells, a step-wise validation of the gating strategy was performed. Detection of GFAP-positive cells was used for n=6 samples where GFAP positivity was validated by staining during neuropathological workup of the same tumors. N=4 patients received 5-aminolevulinic acid (5-ALA) guided surgery and differentiation of tumor cell populations by 5-ALA-positivity in flow cytometry^52^. These n=10 samples were used for cross-validation of autofluorescence-based gating (excitation wavelength 488 nm, emission wavelengths 655-730nm)^53^ which was applied in another n=10 samples without GFAP- or 5-ALA-positivity.

For samples containing very high proportions of TAMs, in the first step, the radioactivity per single TAM cell was calculated using the well purified CD11b(+) TAM enriched fraction (see gating and analysis strategy A in **Extended Fig. 2**). Since the tumor cells in the depleted fraction were not purified to a satisfactory degree (<70%), the radioactivity attributable to the fraction of TAM cells was subtracted from the total activity of the depleted fraction and the remaining radioactivity was divided by the total cell number of tumor cells in the depleted fraction for calculation of radioactivity per single tumor cell. For samples containing very low concentrations of TAM, in the first step, the activity per single tumor cell was calculated in the CD11(−) fraction (see gating and analysis strategy B in **Extended Fig. 2**). Since TAM in the CD11b(+) fractions were not purified to a satisfactory degree (<70%), the radioactivity attributable to tumor cells was subtracted from the measured radioactivity in the TAM enriched fraction and the remaining radioactivity was divided by the total cell number of CD11b(+) cells in the TAM enriched fraction to obtain radioactivity per TAM. Negligibility of TSPO tracer uptake by non-TAM and non-tumor cells for the overall radioactivity in the cell pellets was confirmed by a regression model of cell numbers and gamma emission in TAM enriched fractions of all biopsy samples.

### Small-animal PET/CT

All small animal positron emission tomography (PET) procedures followed an established standardized protocol for radiochemistry, acquisition, and post-processing^54,55^. For tumor and sham mice, [^18^F]GE-180 TSPO small-animal PET (12 ± 1 MBq, n=36) recordings with an emission window of 0-60 minutes after injection were obtained to measure cerebral TSPO expression prior to sorting using a Mediso PET/CT system (Mediso, Budapest, Hungary). A contrast enhanced CT was performed prior to the PET scan. For the mice of the depletion experiment, a static [^18^F]GE-180 emission (13 ± 2 MBq) was recorded between 60 and 90 minutes after injection using a harmonized Siemens Inveon DPET (Siemens, Munich, Germany). All small-animal PET experiments were performed with isoflurane anesthesia (1.5% at time of tracer injection and during imaging; delivery 3.5 L/min).

All analyses were performed by PMOD (V3.5, PMOD Technologies, Basel, Switzerland) using CT (tumor and sham mice) and tracer-specific templates (depletion experiment) for spatial normalization^54^. For tumor mice, a 40-60 minute frame was analyzed and normalization of activity was performed by SUV. The average TSPO-PET SUV was obtained from a spherical volume of interest at the tumor site as the primary PET read-out^16^. Within the tumor volume of interest, we performed a 50-step k means clustering using the PMOD segmentation tool (10 iterations) to delineate individual clusters of tumor tracer uptake in each tumor. For correlation analysis with light-sheet microscopy, a 50% threshold was applied prior to clustering to avoid artificial associations by inclusion of necrotic parts of the tumor.

For the depletion experiment, normalization of injected activity was performed by the previously validated myocardium correction method^56^ and PET estimates deriving from a whole brain volume of interest^54^ were extracted.

### Human PET/CT

TSPO-PET was available for 13 out of 20 (65%) patients that underwent scRadiotracing. All human TSPO-PET scans were performed on a Biograph 64 PET/CT scanner (Siemens, Erlangen, Germany). Tracer production and image acquisition were performed as described previously^57^. Approximately 180 MBq [^18^F]GE-180 were injected as an intravenous bolus and summation images 60-80 minutes p.i. were used for image analysis using a Hermes workstation (Hermes Medical Solutions, Stockholm, Sweden). MRI sequences included gadolinium-enhanced T1- and T2-weighted images and were used for anatomical mapping of stereotactic biopsy coordinates and resected tumor mass within the TSPO-PET images using BRAINLAB ELEMENTS™ (Brainlab AG, Munich, Germany). For correlation analysis between scRadiotracing and TSPO-PET imaging, the mean SUV was obtained from a 0.2 cm³ (25 voxels) volume-of-interest using the sample coordinates.

Amino acid PET was available for a subset of the investigated patients (80%) and served for characterization of vital tumor tissue. Approximately 180 MBq [^18^F]FET were injected and summation images 20-40 minutes p.i. were analyzed. Tumor uptake was assessed as maximum standardized uptake value (SUV_max_). As described previously^58^, the mean background activity was defined as the mean activity of at least 6 crescent-shaped cortical areas in the healthy contralateral side and SUV_mean_ and SUV_max_ were divided by the mean background activity to obtain mean and maximal tumor-to-background ratios (TBR_mean_ and TBR_max_). The biological tumor volume was semiautomatically delineated using the standard 1.6 x background activity as threshold.

### Neuropathological analysis

5 µm thick sections of formalin-fixed and paraffin-embedded (FFPE) tumor tissue were routinely stained with Hematoxylin and eosin (H&E). In addition, immunohistochemical stainings using antibodies against GFAP (polyclonal; Agilent Technologies, Santa Clara, CA, USA), MAP2 (HM-2; Merck, Darmstadt, Germany), IDH 1 (R132H) (QM002; quartett GmbH, Potsdam, Germany), ATRX (BSB-108; Bio SB, Goleta, CA, USA) and Ki67 (MIB1; Agilent Technologies, Santa Clara, CA, USA) were routinely performed according to standard protocols.

### Transcardial perfusion, immunohistochemistry and tissue clearing

Mice that were intended for 3D-histology were i.p. injected with 100 mg/kg ketamine 10% and 10 mg/kg xylazine 2% in 0.9% NaCl. After expiration of the pedal reflex, intracardial perfusion was performed with 0.1 M PBS (10 U/mL, Ratiopharm) for 6 minutes, followed by the administration of 4% paraformaldehyde (PFA) in 0.1 M PBS (pH 7.4) (Morphisto, 11762.01000) for another 6 minutes. Afterwards, the brain was removed and post-fixed by 4% PFA for 6-12 hours at 4°C and washed with 0.1 M PBS and stored in 0.1 M PBS. Samples were subjected to a modified version of vDISCO^44^ and SHANEL^59^ protocols. Samples were decolorized with 25% CUBIC solution for 1 day. An extra permeabilization step was performed using SHANEL reagents: 10% CHAPS with 25% N-Methyldiethanolamine in dH2O at 37°C overnight in order to access the dense glioblastoma.

Then, samples were further permeabilized with vDISCO permeabilization solution with additional 10% 2-Hydroxypropyl-β-cyclodextrin overnight at 37°C. The next day, antibody labeling and boosting step was performed with the addition of CD11b (1:50) (Miltenyi Biotec, 130-110-611) and GFP-Nanobooster LOT 102 (1:500) (Chromotek, gba647n). The samples were incubated in 37°C for 9 days. After labeling, samples were washed with vDISCO washing solution for 3 hours and 3DISCO clearing^60^ with 50-70-90-100-100% tetrahydrofuran (THF) (Sigma, 186562), 1h each, dichloromethane (Sigma, 270997), 30 minutes, was performed. Lastly, samples were placed in BABB (2:1, Benzyl Benzoate, Benzyl alcohol) (Sigma, 24122 and W213802) solution indefinitely for refractive index matching and light sheet fluorescent imaging.

### Light sheet fluorescent microscopy

Ultramicroscope Blaze (Miltenyi Biotec) was used to acquire single plane illumination (light sheet) image stacks. Filters used were ex 470/40 nm, em 535/50 nm; ex 545/25 nm, em 605/70 nm; ex 560/30 nm, em 609/54 nm for autofluorescence, CD11b signal and cancer cells respectively. For all scans, 1.66x magnification, 1×2 tiling, 30% overlap, 120 ms exposure time, 100% light sheet width, ∼7 μm laser sheet thickness, ∼0.31 N/A and 5 μm z-step was used. In order to prevent signal saturation, laser power was adjusted per each sample.

### Cryosectioning and immunofluorescence staining

To determine the absolute cell numbers within tumors, after acquiring the light sheet images of brains, tissue was rehydrated for sectioning. After sequential incubation in THF, samples were washed in PBS and placed in 30% sucrose solution for dehydration overnight. The next morning, samples were embedded in OCT and frozen. 15 μm sagittal sections were acquired using a cryostat (CM3050S; 665 Leica, Wetzlar, Germany). Tissue sections blocked 0.2% TritonX-100, 10% DMSO,10% goat serum in 0.1 M PBS for 1 hour, stained, CD11b (1:50) and GFP-Nanobooster LOT 102 (1:500) in 0.2% Tween-20, 5% DMSO, 5% goat serum, 0.001% heparin in 0.1 M PBS and washed with 0.2% Tween-20, 0.001% heparin in 0.1 M PBS for 3×2 minutes.

### Laser scanning confocal microscopy

Leica SP8 was utilized for laser scanning confocal microscopy. Images were acquired with 40x magnification acquired as 8-bits with HC PL APO CS2 40 × /1.30 NA HC PL APO CS2 40×/1.30 NA or a HC PL APO CORR CS2 63×/1.30 NA objective, 1,024 × 1,024 resolution, 200 Hz, at 20–25°C. Tile scans were obtained with z-step size of 1 μm to allow 3D reconstruction of tissue for depth of 15 μm.

### Image processing

Tiles from each brain were stitched as described previously^44^. Stitched images were saved in TIFF format for further processing. Volumes of segmented CD11b-positivity per TAM and segmented GFP-positivity per tumor cell were obtained via confocal images to allow calculation of absolute cell numbers in 3D-histology (see **Equation 1**). Regions of interest with predominant CD11b (n=5) and cancer cell (n=5) signal were acquired and DAPI nuclear stains were used to count cells per ROI. To segment single nuclei from cancer cells and CD11b(+) cells for estimation of absolute cell numbers, cellpose^46^ was utilized. The percentage of fluorescence(+) nuclei/cells was determined via co-localization of nuclei with adjacent CD11b- and GFP-positivity. Volumes of CD11b- and GFP-positivity were obtained via assessment of segmented areas per channel (50% max. threshold) for each single layer after applying a Gaussian filter of 6mm. To account for positive areas not belonging to captured nuclei (i.e. at the edge of cells) we determined the area of CD11b- and GFP-positivity remote from nuclei (manual threshold visually matching the cell borders). To account for caption of variable profiles in single cells, and we assumed a Gaussian distribution of modelled percentage areas in a fluorescence positive sphere (i.e. cell) with an internal fluorescence negative sphere (i.e. nucleus), calculating a correction factor of 1.044. Areas per cell were then extrapolated to volumes per cell using spherical assumption.

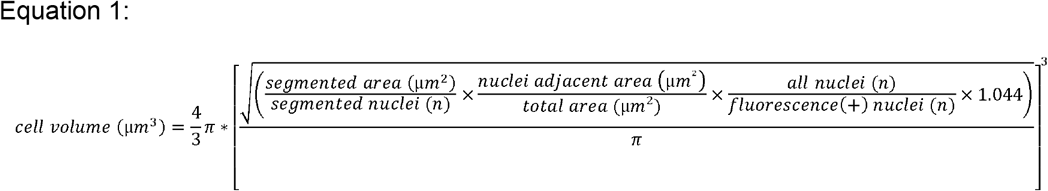

To calculate the volume of tumor cells and TAMs within the tumor in 3D-histology, two channels were separately segmented using syGlass virtual reality tool^45^. The TIFF-stack was imported into the syGlass software where the stack is rendered as a 3D image in a virtual space. The ROI tool was used to annotate the signal of interest with adjustable thresholding in the 3D space to get the best signal to noise ratio. The annotation mask was exported as TIFFs for further downstream processing. Based on the VR-aided manual annotations of tumor and TAM regions, areas were calculated as the sum of the marked mask voxels. LSFM’s resolution of 19.58 mm^3^ per voxel was used to compute the total volume in mm^3^. The percentage of TAMs within the tumor was obtained by first filtering the annotated TAM mask with the tumor area by dot multiplication between the two masks. Then, the number of obtained voxels was divided by the total amount of tumor voxels. Finally, the PET signal of the tumor was compared with the aggregated recovered signals of tumor cells and TAMs (see **Equation 2**).

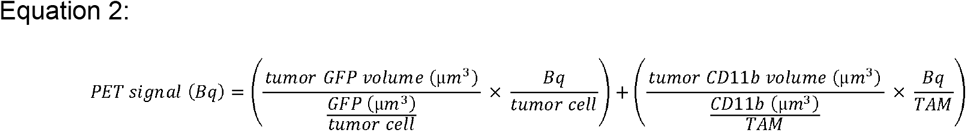

### Proteomics

Isolated TAMs (n=5) and tumor cells (n=3), as well as control microglia isolated from untreated control mice (n=6) were prepared for mass spectrometry based label free protein quantification. The cell pellets were lysed in 50 µl of RIPA lysis buffer (50 mM Tris, 150 mM NaCl, 5 mM EDTA, 1% (v/v) Triton, 0.5% (w/v) sodium deoxycholate, 0.1% (w/v) SDS, pH 8.0) at 4°C with intermediate vortexing. DNA was disrupted by ultrasonication (M220 Focused-ultrasonicator, Covaris, Woburn, MA, USA). The samples were centrifuged for 5 minutes at 16000g and 4°C to remove cell debris and undissolved material. The supernatants were transferred to protein LoBind tubes (Eppendorf, Hamburg, Germany). Proteins were reduced at 37°C for 30 minutes with 15 mM dithiothreitol (DTT) followed by cysteine alkylation with 60 mM iodoacetamide (IAA) for 30 minutes at 20 °C. Excess of IAA was removed by adding DTT. Detergent removal and subsequent digestion with 0.2 µg LysC and 0.2 µg trypsin (Promega, Walldorf, Germany) was performed using the single-pot, solid-phase-enhanced sample preparation as previously described^61^. After vacuum centrifugation, peptides were dissolved in 20 µl 0.1% formic acid. The peptide amount of tumor cell samples was estimated using a fluorescence based protein assay (Qubit Protein assay, Thermo Fisher Scientific, Waltham, MA, USA).

For mass spectrometry, the injected volume was adjusted on the basis of the counted cell numbers to analyze equal cell numbers with a maximum injection amount of 400 ng of peptides. The peptides were separated on a nanoElute nanoHPLC system (Bruker, Bremen, Germany) using a 5 mm trapping column (Thermofisher Scientific, Waltham, MA, USA) and an in-house packed C18 analytical column (15 cm × 75 µm ID, ReproSil-Pur 120 C18-AQ, 1.9 µm, Dr. Maisch GmbH). Peptides were separated with a binary gradient of water and acetonitrile (B) containing 0.1% formic acid at flow rate of 300 nL/min (0 min, 2% B; 2 min, 5% B; 70 min, 24% B; 85 min, 35% B; 90 min, 60% B) and a column temperature of 50°C. The nanoHPLC was online coupled to a TimsTOF pro mass spectrometer (Bruker, Bremen, Germany) with a CaptiveSpray ion source (Bruker, Bremen, Germany). A Data Independent Acquisition Parallel Accumulation–Serial Fragmentation (DIA-PASEF) method for spectrum acquisition. Briefly, ion accumulation and separation using Trapped Ion Mobility Spectrometry (TIMS) was set to a ramp time of 100 ms. One scan cycle included one TIMS full MS scan and with 26 windows with a width of 27 m/z covering a m/z range of 350-1002 m/z. Two windows were recorded per PASEF scan. This resulted in a cycle time of 1.4 s. The data analysis and protein label-free quantification (LFQ) was performed with the software DIA-NN^62^ version 1.8. The data was analyzed using a spectral library including 13912 protein groups and 208218 precursors in 174909 elution groups generated with the same samples and additional murine microglia samples using DIA-NN searching against a one protein per gene database from Mus musculus (download date: 2022-01-25, 21994 entries). Trypsin was defined as protease and 2 missed cleavages were allowed. Oxidation of methionines and acetylation of protein N-termini were defined as variable modifications, whereas carbamidomethylation of cysteines was defined as fixed modification. The precursor and fragment ion m/z ranges were limited from 350 to 1002 and 200 to 1700, respectively. Precursor charge states of 2-4 were considered. The mass accuracy for peptides and peptide fragments was set to 15 and 20 ppm, respectively. A FDR threshold of 1% was applied for peptide and protein identifications. Data normalization was disabled to obtain quantification values, which reflect the protein amounts per cell.

Ratios of TSPO protein expression between SB28 tumor cells, TAMs and control microglia were compared against corresponding ratios obtained by scRadiotracing. Furthermore, all proteins were analyzed regarding their SB28 tumor cell and TAM expression levels relative to control microglia. Potential TAM-specific radiotracer targets were identified in a multi-step process. First, all proteins with higher levels in TAMs (TAM-to-control-microglia ratio) when compared to TSPO were selected. Second, among these, we identified proteins that were not present in SB28 tumor cells or showed high specificity in TAMs over SB28 tumor cells (TAM-to-tumor cell ratios ≥10). Third, to further interrogate specificity, the Human Protein Atlas^47^ was used to analyze single cell RNA expression levels of brain and non-brain cells for all identified target proteins that could serve for subsequent development of TAM-specific radioligands. Fourth, the predominant cell type and the functional immune cell cluster of the identified proteins for radiotracer development was determined by the Human Protein Atlas.

### Statistics

Statistical analyses were performed with Graph Pad Prism (V9, San Diego, CA, USA) and SPSS (V26, IBM, Armonk, NY, USA). A significance threshold of p < 0.05 was considered as significant for all experiments.

*Validation of TSPO scRadiotracing:* Radioactivity per cell was compared between microglia and astrocytes by a paired Student’s t-test. In the depletion experiment, TSPO-PET measures of whole brain were compared between treatment and vehicle by an unpaired Student’s t-test. Iba1 immunoreactivity was compared between treatment and vehicle groups by an unpaired Student’s t-test. The decrease of the TSPO-PET signal in the treatment group was considered as microglia bound radioactivity in brain and compared to extrapolated radioactivity of the microglia cell population as assessed by scRadiotracing using a Student’s t-test.

*scRadiotracing in SB28 glioblastoma mice:* Radioactivity per cell (%ID*BW normalized) was compared between different cell populations (tumor cells, TAMs, sham microglia, control microglia) by a one-way analysis of variance with Tukey post hoc test or by a paired Student’s t-test for populations that were present in the same animals. Multiple regression was used to determine contributions of tumor cells [GFP(+)], microglia [CD11b(+)] and astrocytes [ACSA2(+)] to the radioactivity in the cell pellet. As a validation, measured radioactivity in the depleted fraction of the tumor was correlated with predicted radioactivity by cell count and single cell tracer uptake (Pearson’s coefficient of correlation).

*Association between scRadiotracing and PET imaging:* Lesion site SUV was compared between SB28 and sham mice by an unpaired Student’s t-test. Single cell tracer uptake (tumor cells, TAMs and a summed vector) was correlated with the tumor PET signal and between cell types using Pearson’s coefficient of correlation and we applied a regression model with both cell types as predictors and the PET signal as dependent variable. PET cluster values of the tumor segmentation were correlated with single cell tracer uptake and plotted as a function of cluster grade, after determining the best curve fit (linear, quadratic, exponential) by Akaike information criteria. The difference between the cluster agreement of tumor cells and TAMs was calculated to determine dominant contribution of cell types and likewise plotted as a function of cluster grade.

*Human in vitro scRadiotracing:* A regression model with cell count of TAMs and non-TAM/non-tumor cells as predictors and pellet radioactivity as dependent variable was applied to test for significant contribution of non-TAM/non-tumor cells to the total radioactivity. Single cell tracer uptake was compared between cell types and between patients with high- and low-grade glioma by an unpaired Student’s t-test. Single cell tracer uptake (tumor cells and TAMs) was correlated with the tumor PET signal using Pearson’s coefficient of correlation.

*Integrated model of PET, scRadiotracing and 3D-histology*: Comparisons of GFP(+) and CD11b(+) volumes and cell counts in SB28 tumors were performed using a paired Student’s t-test. The following analysis was performed using the 50 cluster regions of interest per individual mouse. TSPO-PET SUVs were multiplied with the regional correlation coefficient of either tumor or TAM single cell tracer uptake correlations with PET. The individual single cell tracer uptake was estimated according to TSPO-PET SUV of the tumor (**Fig. 3B**) and applied to predict light sheet signal intensity per cluster as a surrogate for cell density. This was performed separately for tumor cells and TAMs. A linear regression was performed to test for the agreement between predicted and standard of truth cell density per region. Finally, a combined tumor cell and TAM model was used to predict absolute radioactivity in SB28 tumors as measured by means of TSPO-PET. To this end, the cell count of both tumor cells and TAMs in seven SB28 tumors was multiplied with the individual single cell tracer uptake (as estimated according to TSPO-PET SUV of the tumor) and aggregated before correlation with equally normalized (%ID*BW) PET radioactivity.

*Proteomics*: The statistical data analysis of the DIA-NN output was performed with the software Perseus Version 1.6.14.0^63^. Only LFQ intensities from protein groups with at least two peptides were considered. The protein LFQ intensities were log2 transformed. A 1-way ANOVA test was applied to determine statistically significant differences between the means of the three groups. Afterwards, individual Student’s t-tests were applied to evaluate proteins with a significantly different abundance between the TAMs, tumor cells and control microglia. Additionally, a permutation based false discovery rate estimation was used with a FDR of 5% at s_0_ = 0.1 as threshold^64^.

All numeric values are reported as average group values ± standard error of the mean (s.e.m.) unless otherwise indicated.

## Supporting information

Proteomics_source_file

## Data availability

All source data (i.e. dynamic PET imaging data) is available per reasonable request to the corresponding author. Proteomics source data is available as extended data.

## Author Contributions

LMB: performed tumor inoculation, performed PET acquisition, image analysis and interpretation of PET scans, established and performed scRadiotracing in mice and human tissue samples, performed scRadiotracing analyses, integrated data of 3D-histology and proteomics, wrote the first draft of the manuscript with input of all co-authors. SVK, SQ and JB: contributed and harvested SB28 cells, supported establishment of scRadiotracing, interpreted PET data in the context of human glioblastoma imaging, supported human scRadiotracing by suggesting patients and performing neurosurgical tissue acquisition. ZIK, SU, LH, IH: performed immunohistological analyses and 3D-histology, interpreted histological data and contributed representative images. SAM: performed and interpreted proteomics. KW, AH, LG, LHK: participated in preclinical PET acquisition, image analysis and interpretation. VCR, JH: neuropathological examination and characterization of the tumor samples. STK, PB, MA, LH, DM: supported scRadiotracing data acquisition and interpretation. AZ: performed 3-dimensional PET image reconstruction. NB: performed immunohistological analyses and interpretation of CSF1R inhibition experiments. SL: performed radiochemistry and TAM target identification. PB, MJR, SZ, JH, SFL, AE, JCT, LvB, NLA and MB: conception and design, contributed to interpreting data, enhancing intellectual content of manuscript. All authors contributed with intellectual content and revised the manuscript.

## Acknowledgments

We thank Rosel Oos and Giovanna Palumbo for excellent technical support during PET imaging.

