## Supplement for "Deciphering sources of PET signals in the tumor microenvironment of glioblastoma at cellular resolution"

**Table of Contents**

**Extended Figures**

**eFigure 1 – Comparison of relative single cell TSPO values of scRadiotracing and Proteomics**

**eFigure 2 – Illustration of gating strategies in human tumor tissue samples**

**eFigure 3 – Comparison of 3D TSPO-PET, predicted 3D-histology and 3D-histology**

**eFigure 4 – Heat-map screening for TAM-specific proteins of interest.**

**eFigure 5 – Protein abundance of TAM-specific proteins of interest**

**Extended Tables**

**Extended Table 1 – Supplemental characteristics of the human glioma cohort**

**Proteomics source file (xls sheet)**

**Extended Figure 1**

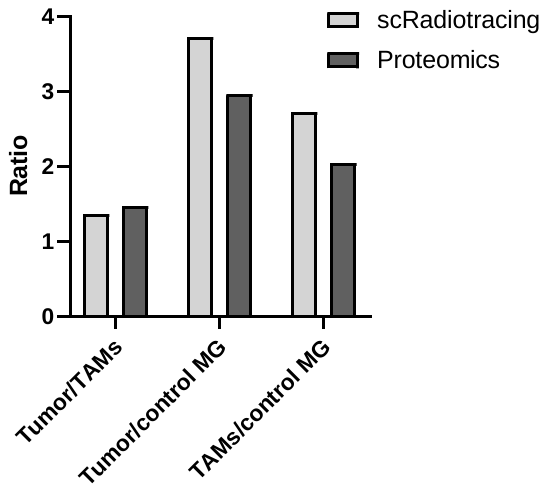

**Extended Figure 1: Comparison of relative single cell TSPO values of scRadiotracing and Proteomics**

Bar graphs show similar relative changes of TSPO levels in single SB28 tumor and tumor-associated microglia/macrophages (TAMs) cells when comparing TSPO tracer uptake as determined by scRadiotracing and TSPO protein levels as determined by proteomics. MG = microglia.

**Extended Figure 2**

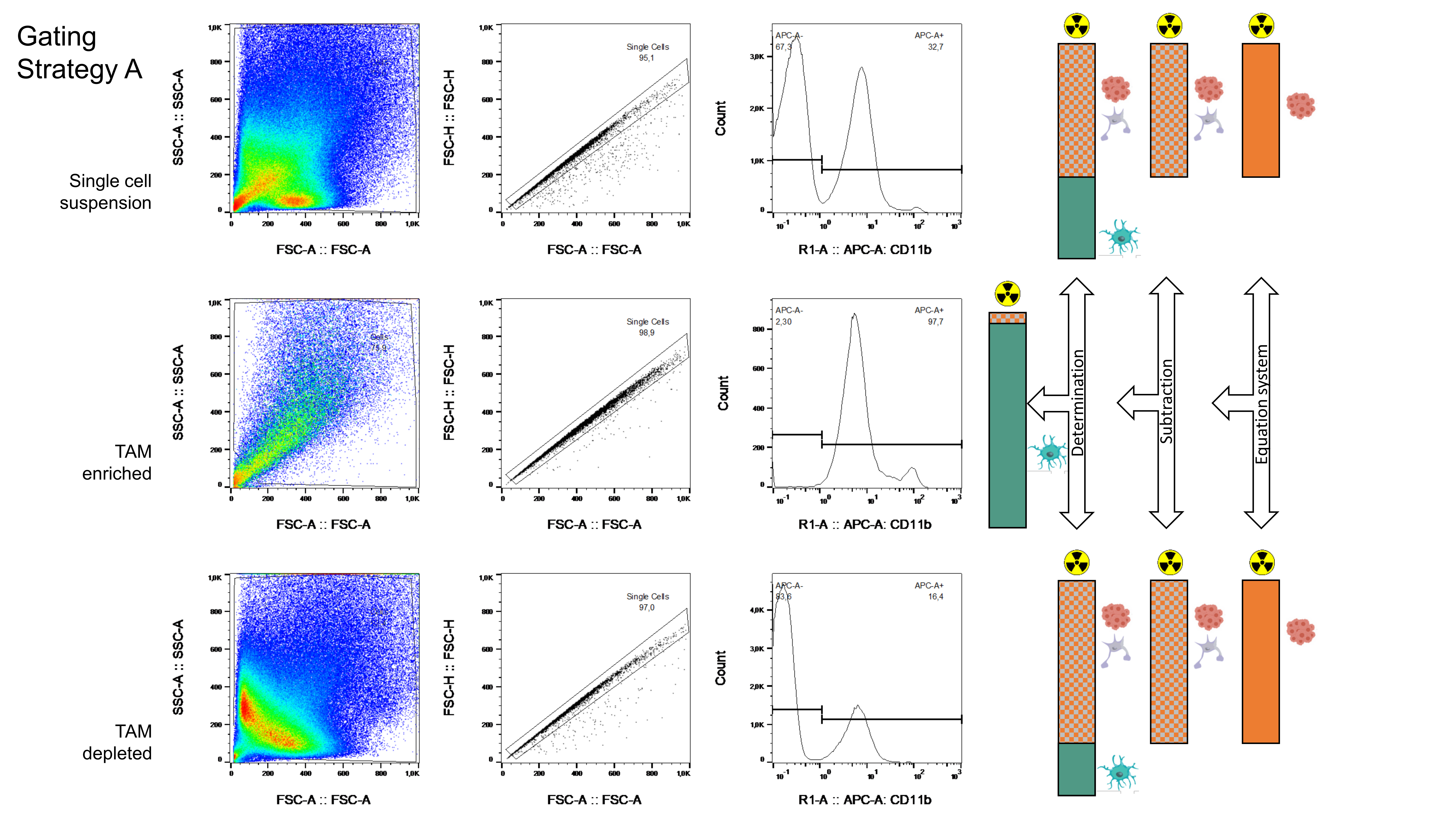

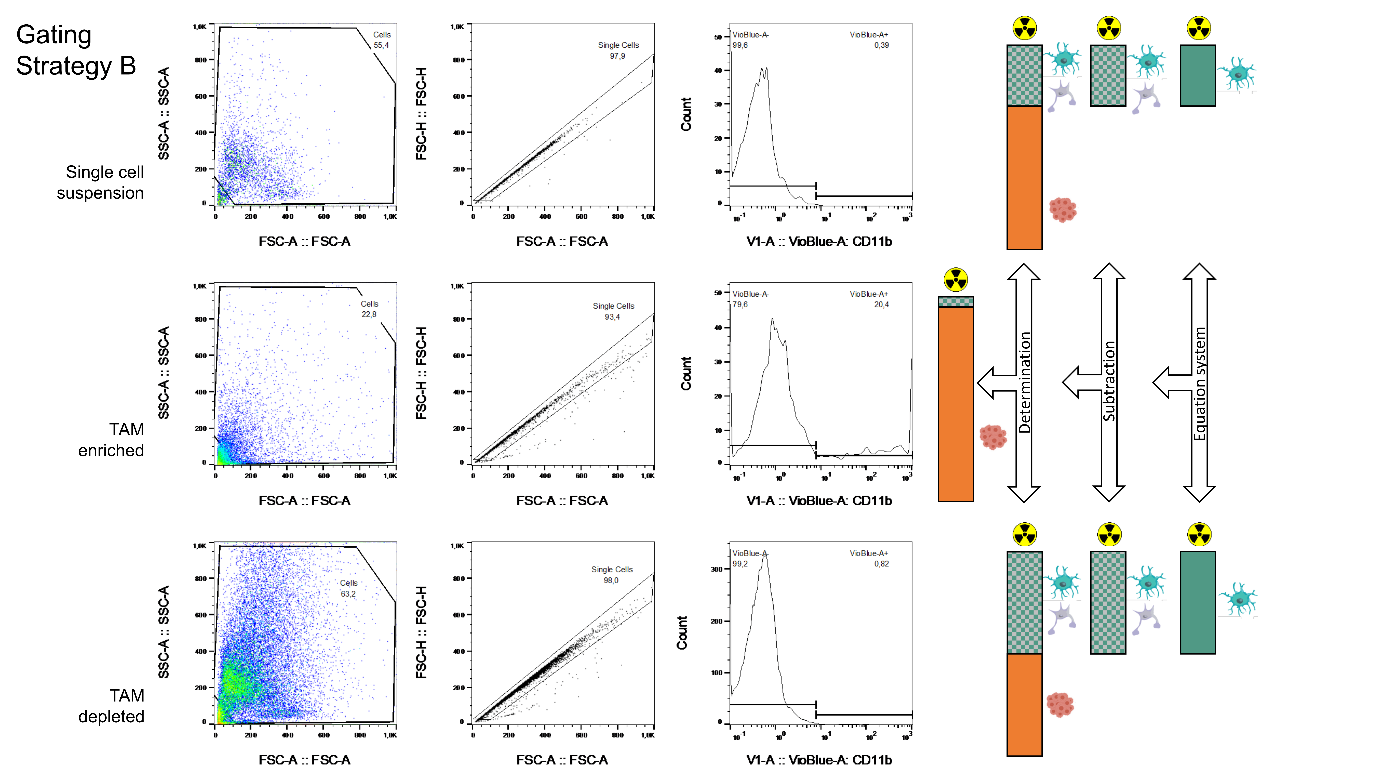

**Extended Figure 2: Illustration of gating strategies in human tumor tissue samples**

For samples containing very high proportions of TAMs, in the first step, the radioactivity per single TAM cell was calculated using the well purified CD11b(+) TAM enriched fraction (see Gating Strategy A). The radioactivity attributable to the fraction of TAMs was subtracted from the total activity of the depleted fraction and the remaining radioactivity was divided by the total cell number of tumor cells in the depleted fraction for calculation of radioactivity per single tumor cell. For samples containing very low concentrations of TAM, in the first step, the activity per single tumor cell was calculated in the CD11(-) fraction (Gating Strategy B). The radioactivity attributable to tumor cells was subtracted from the measured radioactivity in the TAM enriched fraction and the remaining radioactivity was divided by the total cell number of CD11b(+) cells in the TAM enriched fraction to obtain radioactivity per TAM.

**Extended Figure 3**

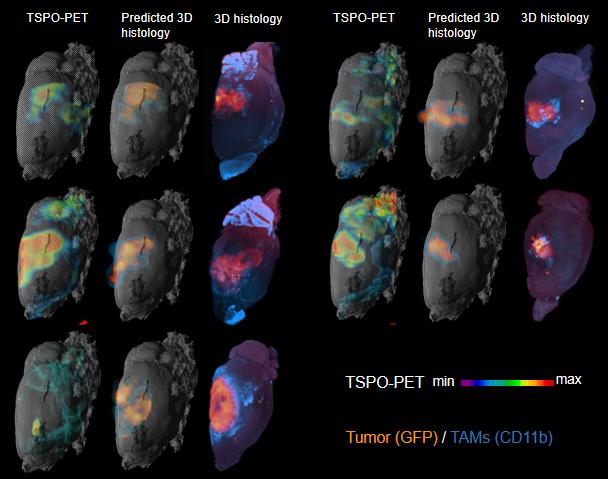
 **Extended Figure 3: Comparison of 3D TSPO-PET, predicted 3D-histology and 3D-histology**

Regional TSPO-PET signals (left) were combined with single cell tracer uptake values of tumor cells and TAMs to predict cell type abundance within individual SB28 tumors. Predicted 3D-histology (middle) shows high regional agreement with standard of truth 3D-histology as obtained by light sheet microscopy (right). Note that even the SB28 tumor with very low signal in TSPO-PET predicted 3D-histology. One sample was imaged and analyzed in the whole brain instead of the tumor hemisphere due to a small lesion near to midline (not shown due to different intensity in 3D visualization).

**Extended Figure 4 (Part A)**

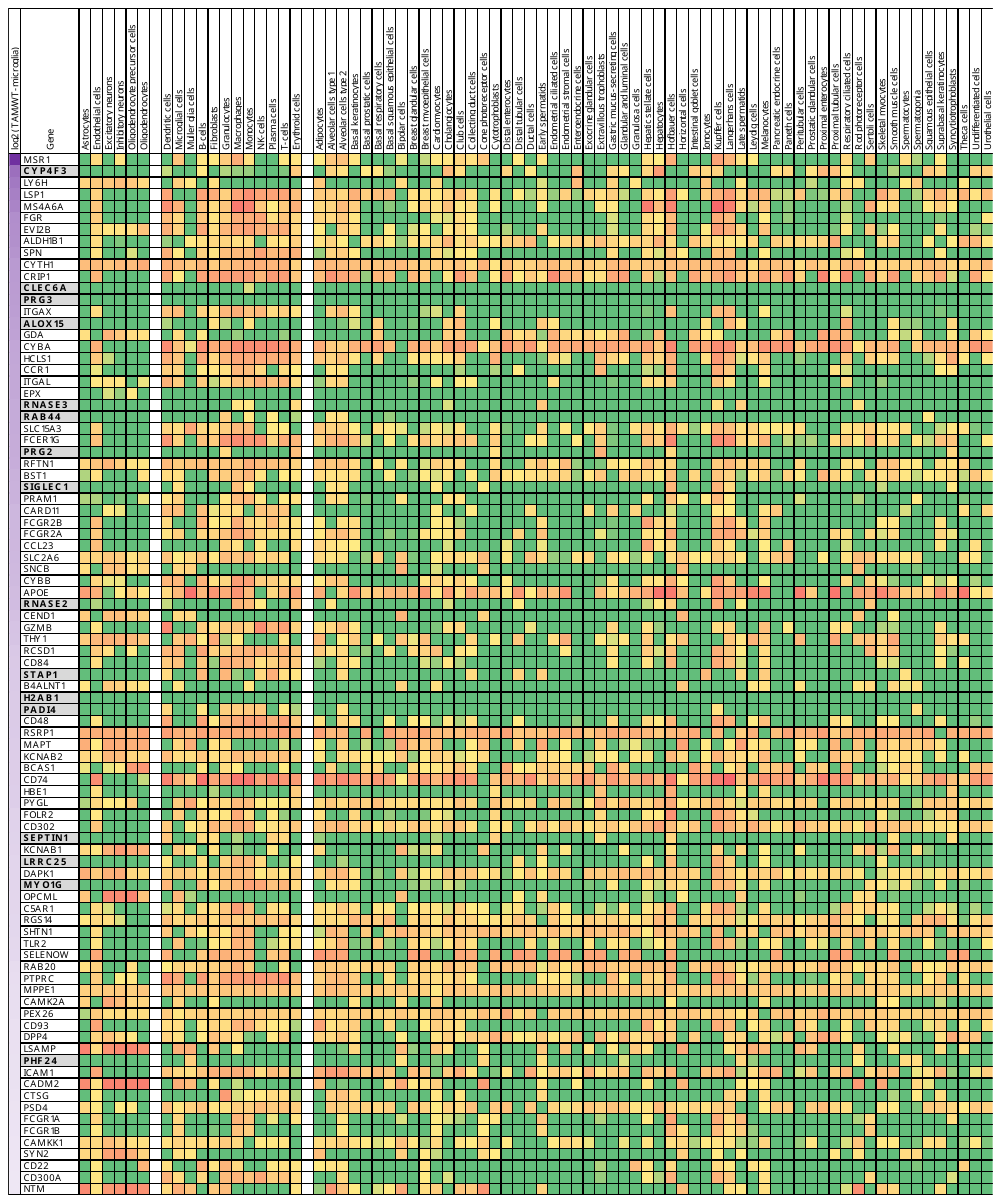

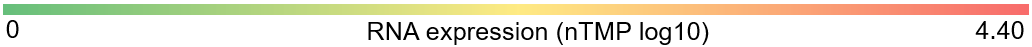

**Extended Figure 4 (Part B)**

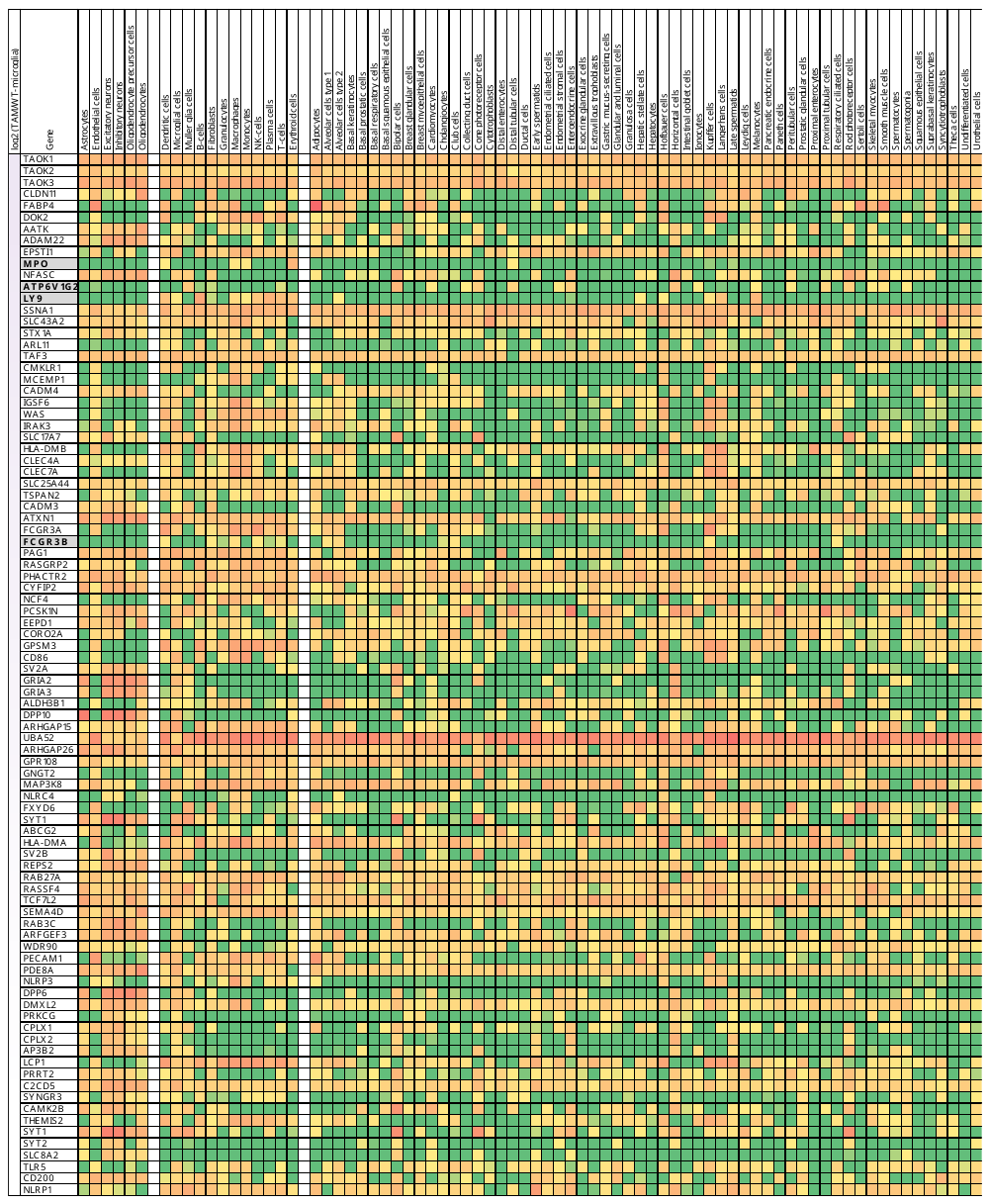

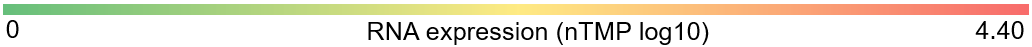

**Extended Figure 4: Heat-map screening for TAM-specific proteins of interest.**

The Human Protein Atlas was used to determine RNA expression levels of all proteins of interest in i) resident off-target cells of the brain (left cell type columns), ii) resident and infiltrating cells in the presence of glioblastoma (middle cell type columns) and iii) off-target cells of the organism (right cell type columns). Genes are sorted by protein level differences in TAMs compared to control microglia of therapy naïve mice (top to bottom; log2 label-free quantification ratio; first column). nTMP = normalized expression.

**Extended Figure 5**

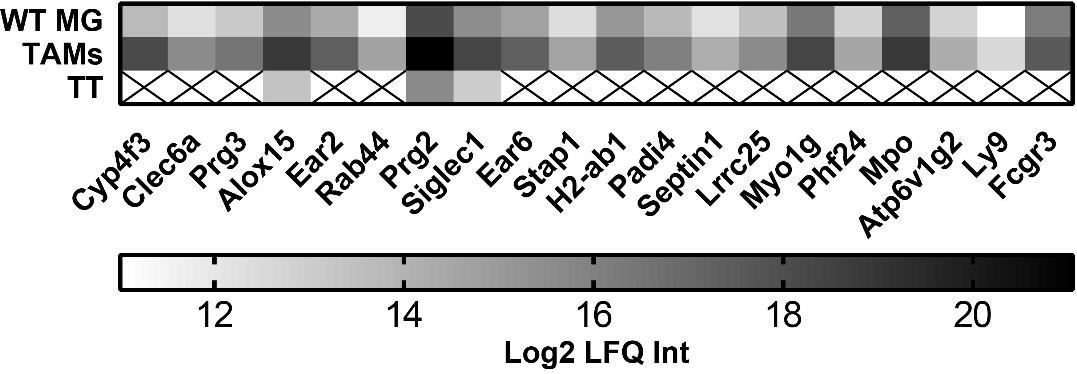

**Extended Figure 5: Protein abundance of TAM-specific proteins of interest**

Heat map shows quantitative protein abundance of normal microglia (WT MG), tumor associated microglia and macrophages (TAMs) and tumor cells (TT) for n=20 TAM-specific proteins of interest. All values are expressed das log2 label-free quantitation intensities (LFQ Int).

**Extended Table 1**

| **ID** | **LOH** | **MGMT** | **TERT mutation** | **Karnofsky** | **Recurrence** | **Previous treatment** | **GFAP staining** | **MRI T2 alteration** | | **MRI contrast enhancement** | | **FET-PET positivity** |
| --- | --- | --- | --- | --- | --- | --- | --- | --- | --- | --- | --- | --- |
| #1 | 0 | 1 | 0 | 90 | 0 | 0 | 1 | 1 | | 0 | | 0 |
| #2 | 1 | 1 | n.a. | 70 | 1 | RCT | 0 | 1 | | 1 | | 1 |
| #3 | n.a. | 0 | 1 | 70 | 1 | RCT, Resection | 1 | 1 | | 1 | | n.a. |
| #4 | 0 | 1 | 1 | 100 | 0 | 0 | 1 | 1 | | 0 | | 1 |
| #5 | 0 | 1 | 1 | 70 | 0 | 0 | 1 | 1 | | 0 | | n.a. |
| #6 | 0 | 1 | 1 | 85 | 1 | RCT, Resection | 1 | 1 | | 1 | | n.a. |
| #7 | n.a. | 1 | 1 | 90 | 1 | RT, Resection | 1 | 1 | | 1 | | 1 |
| #8 | 0 | 1 | 0 | 90 | 1 | CT | 1 | 1 | | 0 | | 1 |
| #9 | 1 | 1 | 1 | 90 | 1 | 0 | 1 | 1 | | 0 | | 1 |
| #10 | 1 | 1 | 1 | 90 | 1 | RCT, Resection | 1 | 1 | | 1 | | 1 |
| #11 | n.a. | 1 | 0 | 90 | 0 | 0 | 1 | 1 | | 1 | | n.a. |
| #12 | 0 | 1 | 0 | 85 | 0 | 0 | 0 | 1 | | 0 | | 0 |
| #13 | 1 | 1 | 0 | 80 | 0 | 0 | 1 | 1 | | 0 | | 0 |
| #14 | n.a. | 1 | 1 | 90 | 0 | 0 | 1 | 1 | | 1 | | 1 |
| #15 | 1 | n.a. | n.a. | 90 | 1 | CT, Resection | 0 | 1 | | 0 | | 1 |
| #16 | 0 | 1 | 0 | 90 | 1 | RCT, Resection | 1 | 1 | | 1 | | 1 |
| #17 | 1 | 1 | 1 | 80 | 1 | CT, Resection RCT/Resection | 0 | 1 | | 0 | | 1 |
| #18 | 1 | 1 | 1 | 90 | 1 | RCT, Resection | 0 | 1 | | 1 | | 1 |
| #19 | 0 | 1 | 0 | 90 | 1 | RCT, Resection | 0 | 1 | | 1 | | 1 |
| #20 | n.a. | 0 | 1 | 80 | 0 | 0 | 0 | 1 | | 1 | | 1 |

**Extended Table 1: Supplemental characteristics of the human glioma cohort.** LOH = Loss of heterozygosity 1p/19q co-deletion, MGMT = Methylation of O6-methylguanine-DNA methyltransferase, TERT = Telomerase reverse transcriptase status, RCT = radiochemotherapy, CT = chemotherapy, RT = radiotherapy, GFAP = glial fibrillary acidic, FET-PET = O-(2-[^18^F]fluoroethyl)-L-tyrosine positron-emission-tomography. 1 = positive, 0 = negative, n.a. = not available
